# Well-Paired-Seq2: High-Throughput and High-Sensitivity Strategy for Characterizing Low RNA-Content Cell/Nucleus Transcriptomes

**DOI:** 10.1101/2023.11.24.568621

**Authors:** Kun Yin, Meijuan Zhao, Yiling Xu, Zhong Zheng, Shanqing Huang, Dianyi Liang, He Dong, Ye Guo, Li Lin, Jia Song, Huimin Zhang, Junhua Zheng, Zhi Zhu, Chaoyong Yang

## Abstract

High-throughput single-cell RNA sequencing (scRNA-seq) is recognized as a powerful technology for disentangling the heterogeneity of cellular states. However, the Poisson-dependent cell capture and low sensitivity in scRNA-seq methods pose challenges for throughput and for samples with low RNA-content. Herein, to address these challenges, we developed Well-Paired-Seq2 (WPS2) based on size-exclusion and locally quasi-static hydrodynamic principles to realize high efficiency of cell utilization, single cell/bead pairing, and cell-free RNA removal. WPS2 exploits molecular crowding effect, tailing activity enhancement in reverse transcription, and homogeneous enzymatic reaction in the initial bead-based amplification to achieve 3116 genes and 8447 transcripts. With an average of ∼20,000 reads per cell, WPS2 detected 1420 more genes and 4864 more transcripts than our previous Well-Paired-Seq. Using WPS2, we overcame the Poisson limit for the capture of both cells and beads and accurately characterized transcriptomes of low RNA-content single cells and nuclei with high sensitivity. WPS2 was further applied to comprehensively profile transcriptomes from frozen clinical samples. We found that clear cell renal cell carcinoma (ccRCC) has a complex microenvironment, and that chromophobe renal cell carcinoma (chRCC) exhibits abundant copy number variations (CNVs). In addition, metanephric adenoma (MA) was characterized at single-cell level for the first time and some potentially specific markers were revealed. With the advantages of high sensitivity, high throughput, and high fidelity, we anticipate that WPS2 will be broadly applicable in basic and clinical research.

## Introduction

Complex cellular systems contain diverse types of cells, and each type can switch among different biological states.^1^ To understand how complex multicellular systems work, it is essential to conduct research on the functionalities and responses of each cell type. Over the past decade, single-cell RNA sequencing (scRNA-seq) has become a powerful tool for uncovering transcriptional profiles and defining cell identities in biological samples. Tang *et al.* first reported the scRNA-seq method in 2009.^2^ Since then, many scRNA-seq technologies have been developed to improve performance, including the yield of the cDNA library, the coverage of a full-length transcriptome, and the sensitivity of gene detection, such as Smart-seq^3^/Smart-seq2^4^/Smart-seq3^5^, CEL-Seq^6^/CEL-Seq2^7^, and Quartz-Seq^8^/Quartz-Seq2^9^. Although these methods have achieved superior performance, the single-cell libraries still need to be constructed individually, which is limited to analyzing tens to hundreds of single-cell transcriptomes at a time. The low throughput and high cost of these scRNA-seq methods make it difficult to comprehensively dissect broadly varied cell types and states of complex cellular systems.^10^

Throughput is an important feature for the deconvolution of cell type and state in complex multicellular systems.^11^ Recently, several high-throughput bead-based methods have been reported.^10, 12, 13^ The throughput has been successfully expanded from a handful of cells to thousands of cells per assay. This increase in cell throughput has promoted the efforts to profile the atlas of whole organs or entire organisms.^13, 14^ However, Poisson-dependent single-cell isolation leads to many unused reactors, which limits the throughput of scRNA-seq due to reagent costs and the physical constraints of devices.^11, 15, 16^ Recently, an integrated dielectrophoresis (DEP)-trapping nanowell-transfer (dTNT-seq) platform was reported for high throughput scRNA-seq, overcoming Poisson-dependent single cell/bead isolation.^16^ However, dTNT-seq requires DEP assistance and precision control of flow rate for single-cell isolation, leading to complicated chip fabrication and long single-cell isolation time. This may result in non-ideal cell viability during single-cell isolation and limit the expansibility of throughput.

To address these limitations, we developed a size-exclusion and locally quasi-static hydrodynamic microwell-based Well-Paired-Seq (WPS) platform^15^, which shows excellent efficiency of single-cell isolation within a short time and without peripheral specialized instrumentation. Compared to the other well-based methods, WPS allowed ∼80% of single-cell/bead pairing, which greatly enhanced the isolation density of single cells and the throughput. Using WPS, we have successfully realized ∼100,000 single-cell analyses per flow channel. Moreover, the presence of cell-free RNA and aggregated cells resulting from tissue digestion usually poses a significant challenge in scRNA-seq. WPS has demonstrated a high degree of effectiveness in removing these undesirable interfering substances, resulting in significantly reduced background noise and a lower risk of generating inaccurate biological findings.^17^ However, WPS still suffered from low sensitivity of gene detection. The inefficiencies in gene detection limited the ability to characterize essential but typically sparsely expressed genes, such as transcription factors, signaling molecules, and affinity receptors, and to reveal distinct cell states, particularly for low-RNA content units, such as immune cells and nuclei.

In addition, the current requirement of harsh enzymatic dissociation for preparing single-cell suspensions from fresh tissues poses a significant obstacle for handling clinical materials, frozen samples, and tissues that cannot be readily dissociated.^18^ Furthermore, single cells that have been treated with enzymes often pose difficulties, such as damage in complexity of RNA molecules by enzymes, skewed proportions of dissociated cell types, and triggered stress reactions of transcriptional expression.^19^ As a complementary technology to scRNA-seq, single-nucleus RNA sequencing (snRNA-seq) can handle complex tissues that cannot be dissociated, thus providing access to archived samples.^18–20^ To date, the development of a compatible method for single-cell and single-nucleus transcriptome profiling with high sensitivity, throughput, and fidelity remains challenging.

Herein, we developed an optimized method, Well-Paired-Seq2 (WPS2), to overcome the limitation of low sensitivity, while being fully compatible with both single-cell and single-nucleus RNA-seq. By adopting molecular crowding effect, tailing activity enhancement in reverse transcription, and homogeneous enzymatic reaction in the initial bead-based amplification, the sensitivity of WPS2 is highly enhanced. After optimization, we were able to detect 3116 genes and 8447 transcripts of NIH 3T3 cells at an average of ∼20,000 reads per cell, with an improvement of 1420 genes and 4864 transcripts compared to our previous WPS. To further validate the performance of WPS2, we applied it to low-RNA content samples, such as immune cells and nucleus samples. We detected an average of 1345 genes and 3180 transcripts at an average of ∼23,000 reads per cell using mouse spleen tissue. This was 680 and 1969 more genes and transcripts than WPS. Using the nuclei from the NIH 3T3 cells and the frozen kidney tissue, we validated the compatibility of WPS2 with snRNA-seq. The successful application to snRNA enables access to massively archived samples.

Finally, we applied our method to analyze renal tumor samples (clear cell renal cell carcinoma (ccRCC), chromophobe renal cell carcinoma (chRCC), and metanephric adenoma (MA)), and revealed a comprehensive profile of multiple pathologic transcriptomes. We found that ccRCC cells have complex microenvironment and that highly expressed genes are associated with immune response, hypoxia, angiogenesis, etc., while chRCC has more copy number variations (CNVs) in its genome, indicating the instability of the genome. We also characterized MA, a rare benign tumor that accounts for ∼0.2% of adult renal epithelial neoplasms^21^, at the single-cell level for the first time. MA are often misdiagnosed due to lack of specificity in clinical presentation and imaging features. Here, we identified 10 candidate specific markers of MA. We expect WPS2, a high-sensitivity, high-throughput, and high-fidelity platform compatible with scRNA and snRNA-seq, will have broad application in cell biology, precision medicine, and reproductive biology.

## Results

### Workflow of Well-Paired-Seq2

To enable high-throughput and highly sensitive sequencing of single cells or single nuclei, we developed WPS2, which was systematically optimized based on our previously reported size-exclusion and locally quasi-static hydrodynamic microwell-based single-cell RNA sequencing platform (WPS)^15^. The workflow of WPS2 is depicted in **Figure 1**, which includes the following major steps: (1) Cells and barcoded beads are successively trapped in the cell-capture-wells and bead-capture-wells to realize single-cell/bead pairing; before loading barcoded beads, cell-free RNA can be removed by washing buffer. Then, cells are lysed by the dissolved surfactant molecules from the settled surfactant aggregates in the sealing oil and the released mRNA molecules are captured by the barcoded beads with oligo(dT) sequence. (2) The captured RNA molecules on the barcoded beads are reverse transcribed using Maxima RTase with the addition of GTP and PEG. After reverse transcription, the single-strand cDNAs on the barcoded beads are used as templates for the second-strand cDNA synthesis. (3) The double-strand cDNAs are amplified by PCR. (4) The cDNA products are used for library preparation using Tn5 transposase. After sequencing, the transcriptome of single cells is inferred from the digital expression matrices and used for downstream analysis.

**Figure 1.**
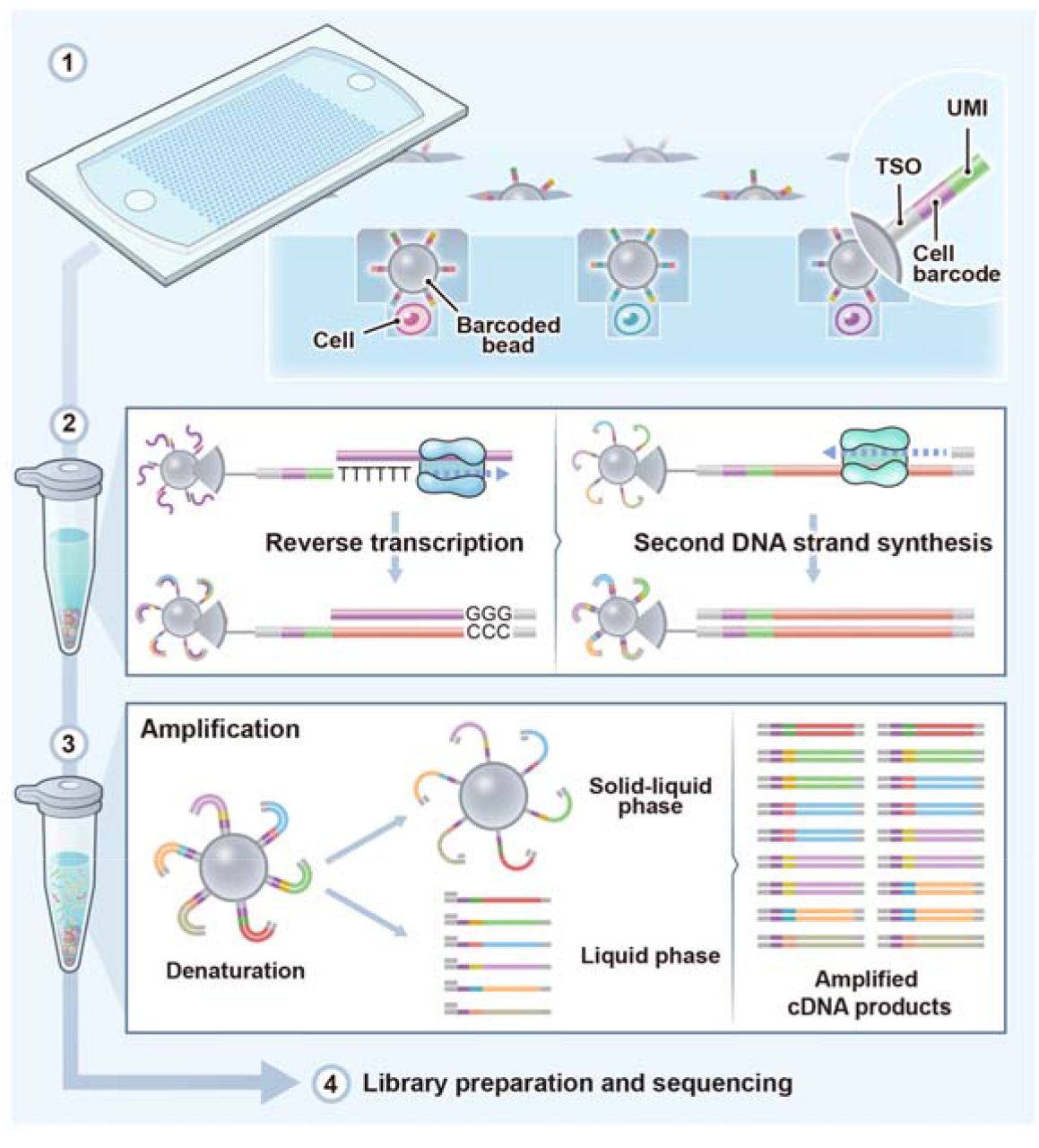
Workflow of Well-Paired-Seq2: (1) Pairing cells and barcoded beads in the size-exclusion and locally quasi-static hydrodynamic dual wells; (2) Reverse transcription of the captured mRNA molecules on the barcoded beads, followed by second strand synthesis; (3) Denaturing the double-strand cDNA molecules and amplification by PCR; (4) Construction of the library of the amplified cDNA products for sequencing.

### Optimizations of WPS2

WPS2 involves several major steps including mRNA capture, reverse transcription (RT), cDNA amplification, and library construction. To improve the quality of scRNA-seq, we tested various parameters, such as incubation time and additives of hybridization buffer in mRNA capture, the time and additives of reverse transcription (RT) buffer in RT, the double-strand cDNA as the initial template for cDNA amplification, and the tagmentation in library construction (**Figure 2, 3 and Figure S1**, S2). We found that the supplementation of additives in RT and double-strand cDNA molecules as the initial templates for PCR significantly improved the gene detection ability. In the RT step, spiking template-switching oligonucleotide (TSO) is commonly used to add a 3’ priming site for whole-transcriptome amplification.^3^ However, the inefficiency of the template-switching reaction compromises the sensitivity of gene detection.

**Figure 2.**
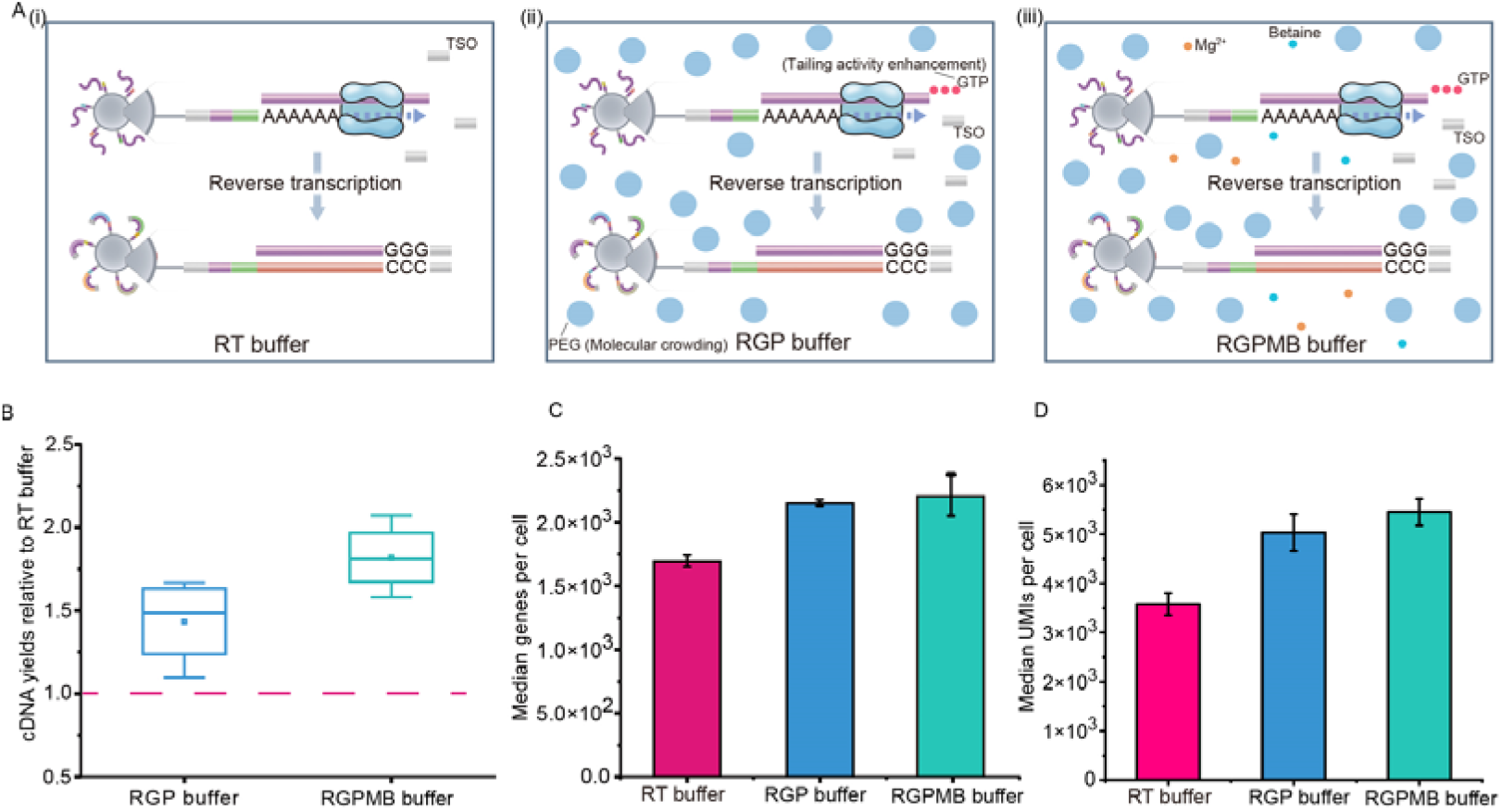
Buffer optimization in reverse transcription. A) Diagram of different additives in RT buffer. B) cDNA yields of RGP buffer and RGPMB buffer relative to RT buffer. C, D) Gene and UMIs detection with three different buffers at an average of 20,000 reads per cell.

To increase the efficiency of the template-switching reaction, several additives were selected as supplements to the RT buffer, including GTP (to enhance the tailing activity)^22^, PEG-8000 (to improve yields by molecular crowding)^23^, Mg^2+4^, and betaine (to improve yields by methyl group donor betaine in combination with high Mg^2+^ concentrations) ^4^. As shown in **Figure 2A**, three types of reaction buffers were tested: RT buffer (Maxima H Minus RT buffer), RGP buffer (Maxima H Minus RT buffer, GTP, PEG), and RGPMB buffer (Maxima H Minus RT buffer, GTP, PEG, Mg^2+^, betaine). **Figure 2B** shows that the cDNA yields of both RGPMB buffer and RGP buffer are significantly higher than that of RT buffer with RGPMB buffer having the highest yield, indicating a higher efficiency of template-switching reaction with RGPMB buffer and RGP buffer. To determine whether the improved cDNA yields indeed correspond to improved sensitivity, we further evaluated the sensitivity of gene detection with these three buffers by sequencing (**Figure 2C, D**). The results showed that, compared to the RT buffer, both RGP and RGPMB buffers led to the detection of more genes and transcripts (medians of 2149 genes and 5030 transcripts for RGP, and 2207 genes and 5449 transcripts for RGPMB, compared to 1696 genes and 3583 transcripts for RT), with an average sequencing depth of 20,000 reads per cell. However, no significant difference was observed in gene numbers between RGP and RGPMB buffer, which is not consistent with the cDNA yields. We speculated that byproducts may be generated when using RGPMB buffer as the RT reaction mix (**Figure S3**). Therefore, the RGP buffer was chosen for the subsequent experiments.

However, this process often results in amplification bias, where low-copy or difficult-to-amplify cDNA molecules are undetectable, limiting the sensitivity of gene detection.^24^ This is an especially critical problem in bead-based scRNA-seq methods. In the process of bead-based scRNA-seq, the first-strand cDNA molecules are covalently attached to the barcoded beads after reverse transcription. Hence, the initial amplification is performed on the solid-liquid interface (heterogeneous reaction) and would impede the efficiency of the second-strand synthesis (**Figure 3A (i)**). Due to the inefficiency of the reaction on the solid-liquid phase, we assumed that only a part of the second strands are synthesized in the initial cycles. However, in subsequent cycles, the synthesized second strands would be released in the denaturation step, and the bias would be largely amplified because of the different efficiencies of strand synthesis at the solid-liquid interfaces and in the liquid phase.

**Figure 3.**
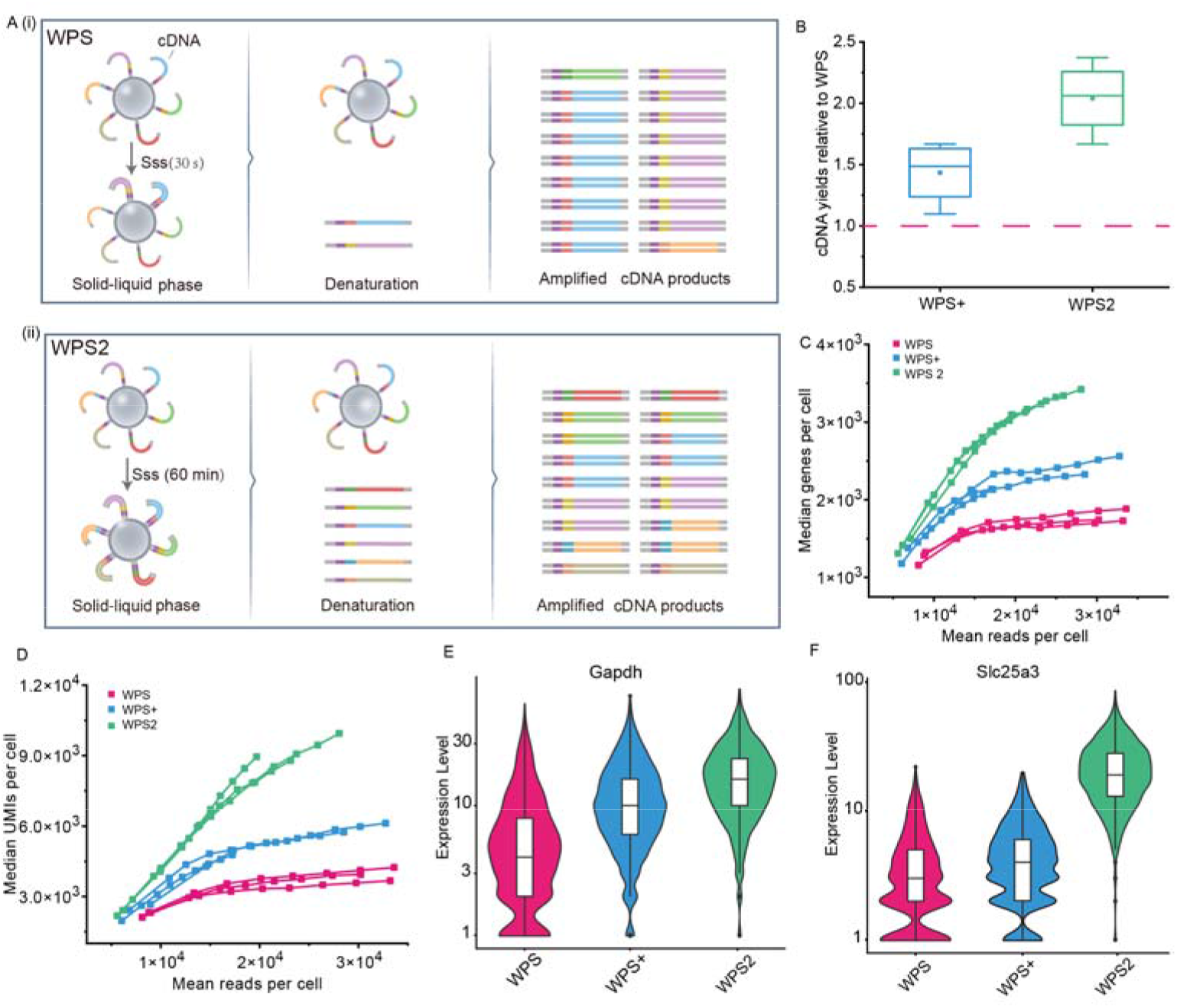
Performance validation of WPS2. A) Diagram of amplification in WPS and WPS2. B) Box plot showing the improvement of cDNA yields relative to WPS. C, D) Median gene and UMI variations along with different mean reads per cell. E, F) Violin plots showing the expression level of given genes (housekeeping genes: *Gapdh*; high variation genes: *Slc25a3*) detected by WPS, WPS+, and WPS2.

To overcome the amplification bias, we designed a second-strand synthesis method (**Figure 3A(ii)**). Before PCR, a long time (60 min) is allowed for second-strand synthesis to ensure that as many as possible second-strand cDNA molecules are synthesized. Therefore, in the first cycle of amplification, the second-strand cDNA molecules are simultaneously released into liquid during denaturation and amplified in the liquid phase, avoiding the efficiency variance of amplification between the solid-liquid interface and the liquid phase.

To assess the sensitivity after adding the second-strand synthesis step, we compared the gene detection of the following three methods: WPS, WPS with RGP buffer (WPS+), and WPS with the RGP buffer and the second-strand synthesis (WPS2). **Figure 3B** shows that the cDNA yields with WPS2 are significantly higher than the yields with WPS (2-fold) and WPS+ (1.4-fold), suggesting a high and uniform efficiency of amplification using WPS2. As expected, the detected genes and transcripts by WPS2 are significantly higher than that of WPS and WPS+ (**Figure 3C, D**). At an average of 20,000 reads per cell, a median of 3112 genes and 8485 transcripts were detected among NIH 3T3 cells in WPS2. Compared to WPS, WPS2 identified 1416 and 4902 more genes and transcripts (**Figure S4A, B**). Zooming in on four genes (two housekeeping genes and two high variation genes in NIH 3T3), we found that WPS2 detected more molecules per cell than WPS and WPS+ (**Figure 3E, F, Figure S4C, D**). Together, these results demonstrated that WPS2 significantly improved the sensitivity after systematically optimizing the workflow, especially the steps of RT and second-strand synthesis.

### WPS2 for low-RNA content sample analysis

To demonstrate the improved performance of WPS2 in profiling low-RNA content samples, we applied the method to characterize the immune cells from spleen tissues. Immune cells and nuclei are often small-sized and low in RNA content, making their isolation and profiling challenging.^25^ In the previously reported WPS method, the cell-capture-well, with a width and height of 25 µm, increased the likelihood of trapping doublets of small cells during the process of cell capture. To overcome this limitation, a smaller cell-capture-well was designed with a width and height of 12 µm, respectively, while the bead-capture-well was designed with an inscribed circle diameter and height of 50 µm (**Figure S5A**). To test the efficiency of the modified dual-well chip for small single-cell isolation, spleen cells (Calcein AM-stained) were loaded and achieved approximately 73.6% of single cells, 1.1% of doublet cells, and 25.3% of zero cells in a 10,000 dual-well chip (**Figure S5B**), which was significantly higher than that of the Poisson distribution-based method (1%-10%)^26^.

After sequencing and filtering, 5029 cells and 10 clusters were identified. Among these clusters, Cd4 T cells were enriched in *Il7r*, *Cd4*, and *Trac*, while Cd8 T cells were characterized by high expression of *Il7r*, *Cd8a*. NK cells were identified by the marker genes of *Nkg7* and *Klre1*, and B cells were defined by the expression of *Ms4a1* and *Cd79a*. Monocyte progenitors were identified by the enrichment of *Ms4a3*, *Elane*, and *Cebpe*, while monocytes were defined by the enrichment of *Cd14* and *Hp*. Macrophage cells were characterized by the high expression of *C1qa*, *C1qb*, and *Vcam1*, and dendritic cells showed the enrichment of *S1100a4*, *Cst3*, and *Itgax*. Erythroblast cells were characterized by the enrichment of *Spc24*, *Cdca5*, *Stmn1*, and *Nuf2*, and plasma cells showed high expression of *Derl3*, *Pon3*, *Prg2*, and *Sdc1*. The spleen was determined to be composed of 31.9% B cells, 18.3% monocytes, 21.6% macrophage cells, 11.2% Cd8 T cells, 6% NK cells, 3.6% dendritic cells, 3.5% Cd4 T cells, 2% erythroblast cells, 1.4% plasma cells, and 0.6 % monocyte progenitors as the rarest of the observed clusters (**Figure 4A-C**). WPS2 detected an average of 1345 genes and 3180 transcripts at an average of ∼23,000 reads per cell. This was 680 and 1969 more genes and transcripts, respectively, than WPS, demonstrating the superior sensitivity of WPS2 (**Figure S5C**). Zooming in on each cell type, we characterized the gene and transcript detection in each cell type (**Figure 4D, E**). In summary, the combination of WPS2 and the modified dual-well chip allows comprehensive cellular deconvolution of cell-type composition in complex multicellular systems with high efficiency of single-cell isolation and superior gene detection.

**Figure 4.**
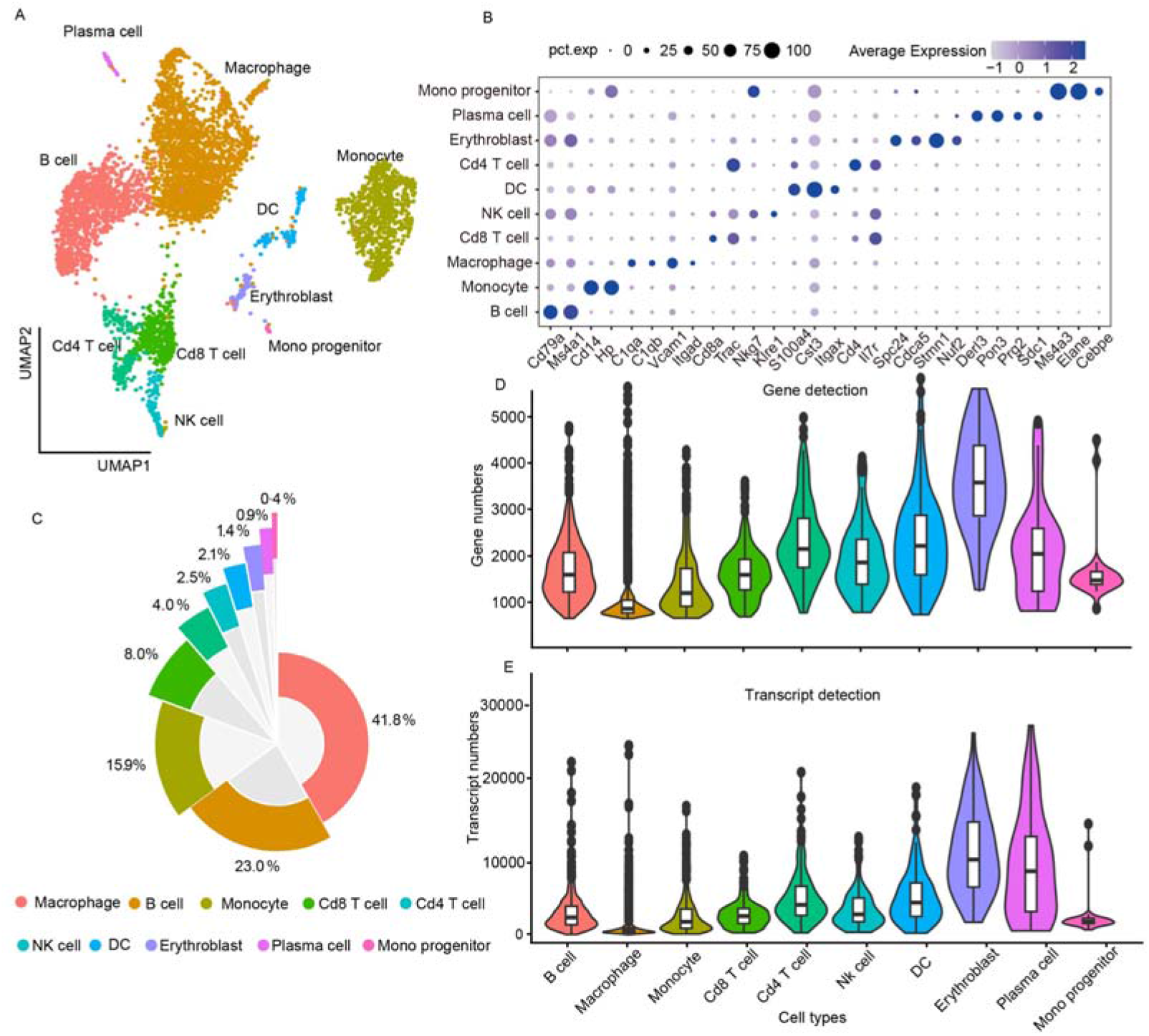
Single-cell transcriptional profiling of mouse spleen cells. A) Unsupervised clustering identifies 10 cell types. Mono progenitor, monocyte progenitor; Plasma cell; Erythroblast; Cd4 T cell; DC, Dendritic cell; NK cell, Natural killer cell; Cd8 T cell; Macrophage; Monocyte; B cell. B) Dot plot visualization of each cell type in spleen by single-cell transcriptome data. C) Pie chart showing the proportion of different cells in the mouse spleen. D, E) Violin plots showing the mean gene and transcript detections across the 10 cell types. 1613, 900, 1306, 1548, 2265, 1884, 2300, 3778, 2058, and 1676 genes, and 3663, 1763, 3312, 3493, 5764, 4414, 6481, 14348,10230, and 3724 transcripts were detected in B cells, macrophage cells, monocytes, Cd8 T cell, Cd4 T cells, NK cells, DC cells, erythroblast cells, plasma cells, and monocyte progenitors, respectively.

### WPS2 for single-nucleus sample analysis

Although scRNA-seq has revolutionized cell biology because of its superior ability to disentangle heterogeneous transcriptional profiles at an unprecedented resolution, the preparation for single-cell suspensions from fresh tissue is a major roadblock to assessing clinical samples, frozen materials, and tissues that cannot be readily dissociated. To tackle these constraints, snRNA-seq was developed to handle samples of complex tissues that are not readily dissociated. To demonstrate that WPS2 is compatible with snRNA-seq, nuclei extracted from NIH 3T3 cells were tested (**Figure S6A**). As shown in **Figure 5A**, the gene numbers of each cell along with read depth were observed. In addition, the average expression in nuclei showed good correlation with the scRNA-seq data (r=0.80, **Figure 5B**), demonstrating the accuracy of WPS2 for snRNA-seq.

**Figure 5.**
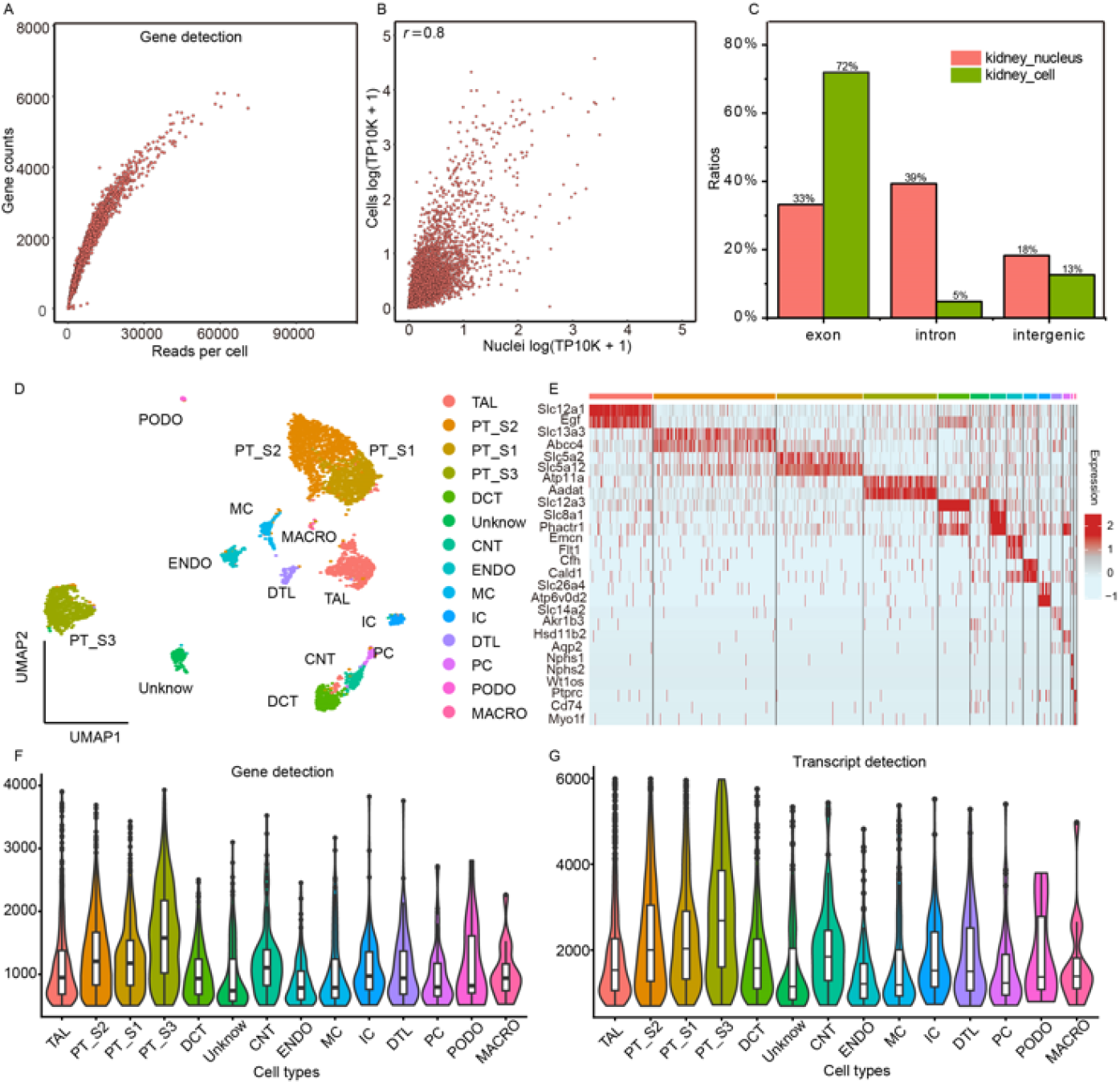
WPS2 for high-throughput single-nucleus RNA-seq. A) Scatter plot showing the gene numbers versus read numbers of each individual NIH 3T3 cell. B) Scatter plot showing Pearson correlation between NIH 3T3 nuclei (x axis) and cells (y axis) by WPS2. Log (TP10k + 1) corresponds to log-transformed UMIs per 10k. C) Percent reads mapped to the exons, introns, and intergenic regions on mouse genome for cells (green bars) from mouse fresh kidney and nuclei (red bars) from mouse frozen kidney. D) Visualization by UMAP plot of clustering of 13 cell-type expression profiles from mouse frozen kidney. TAL (thick ascending limb of Henle’s loop cell), PT_S1 (proximal tubule segment 1), PT_S2 (proximal tubule segment 2), PT_S3 (proximal tubule segment 3), DCT (distal convoluted tubule cell), CNT (connecting tubule cell), ENDO (endothelial cell), MC (mesangial cell), IC (intercalated cell), DTL (descending thin limb of Henle’s loop cell), PC (principal cell), PODO (podocyte), MACRO (macrophage cell). E) Gene expression heatmap showing the top differentially expressed genes for each cell cluster in mouse single-cell data. F, G) Violin plots showing the detection of genes and distribution of the number of transcripts in each cluster.

Next, we applied WPS2 to characterize the nuclei from frozen kidney tissue (**Figure S6B**), resulting in the identification of 4217 nuclei with 1309 mean genes, 2329 mean transcripts, and an average of 55,875 reads per cell (**Table S1**). Moreover, we verified that the data of scRNA-seq and snRNA-seq can be well integrated (**Figure S7**). Since the most nascent RNA molecules are found in the nucleus, while the mature RNA molecules are in the cytoplasm, snRNA-seq can map many reads to introns. Using WPS2, we found that 33% of reads were mapped to introns and 39% were mapped to exons with snRNA-seq, while with scRNA-seq, 5% of these genomic reads were mapped to introns and 72% were mapped to introns (**Figure 5C**).

We then used Seurat to interrogate the cell types and visualize the results in the uniform manifold approximation and projection (UMAP) space. Seurat identified 13 cell types, and specific markers of each cell type were characterized (**Figure 5D, E**). Next, we comprehensively characterized the genes and transcripts of each cell type (**Figure 5F, G, Table S1**). These results verified that WPS2 enables cell-type deconvolution by clustering snRNA-seq profiles and holds great promise to explore the variety of precious samples in the frozen tissue bank.

### WPS2 for clinical sample analysis

Renal cell carcinomas (RCCs) are increasingly common malignancies worldwide, ranking among the top ten most diagnosed.^27^ Despite limited understanding of tumorigenesis for the heterogeneous subtypes of RCC, advanced single-cell technologies provide clinical practitioners with a detailed and novel high-resolution insight into the mechanisms^28^. To better understand the heterogeneity different pathological types in RCC’s and to include a benign tumor for comparison, we employed WPS2 to characterize multiple renal tumors, including ccRCC, chRCC, and MA. In total, we processed 5 biopsies using scRNA-seq (2 samples) or snRNA-seq (3 samples). First, we integrated all the tumor samples, summing to 15,634 cells, and divided them into 11 clusters after data quality filtering. The tumor cell clusters were determined based on the cell type-specific markers. The neoplastic cells representing ccRCC in our cohort overexpressed classic biomarkers of this disease, like CA9 and ANGPTL4; chRCC tumor epithelia overexpressed *KIT* and *RHCG*; and MA overexpressed *WT* and *THNT3* (**Figure 6A, B**). Next, we inferred the copy number variations (CNVs) and Gene Ontology (GO) analyses in three subtypes of RCCs. Chromosome 3p was lost in ccRCC samples (**Figure 6C**), as well as enrichment of hypoxic signaling and vascular transport (**Figure 6E**), considering that the pathogenesis of ccRCC is related to hypoxia caused by *VHL* mutation.^29^ Loss of heterozygosity of chromosomes 1, 2, 6, 10, 13, and 17 is a signature event of chRCC (**Figure 6C**), consistent with previous studies.^29, 30^ The CNV scores in **Figure 6D** show that chRCC has a high degree of chromosomal instability. The differential gene expression patterns enriched in cell growth and expansion suggest that chRCC may have a higher proliferative capacity compared to the other two tumor samples (**Figure 6F**), which is consistent with its aggressive clinical behavior.^30, 31^

**Figure 6.**
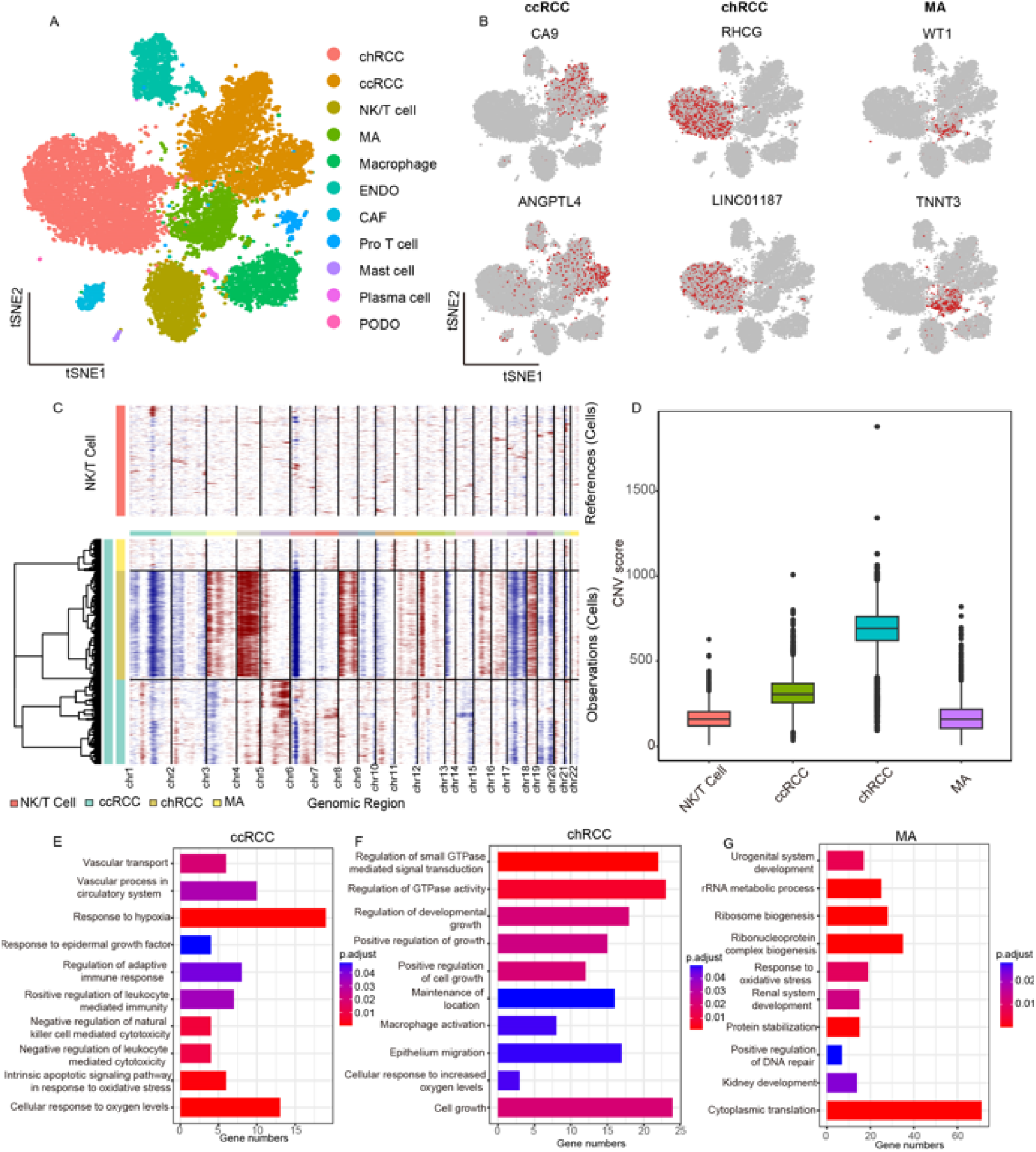
Single-cell transcriptional profiling of clinical renal tumors. A) Visualization by tSNE plot of clustering of 15,634 single-cell and single-nucleus expression profiles (n=5) from clinical renal tumors(n=3). ccRCC, clear cell renal cell carcinoma; chRCC, chromophobe renal cell carcinoma; MA, metanephric adenoma cell; Pro T cell, proliferative T Cell; ENDO, endothelial cell; CAF, cancer-associated fibroblast; PODO, podocyte. B) tSNE plots showing the expression of marker genes for renal tumor cells. C) Clustering of CNV profiles inferred from all sample sequencing data. Heatmap of CNV signal normalized against the NK/T cells shows CNVs by chromosome (columns) for individual cells (rows). D) Comparison of the CNV score across non-malignant cells and RCC cells. E-G) Biological process analysis of gene ontology enrichment for three subtypes of renal tumors.

MA is a rare benign renal neoplasm that accounts for ∼0.2% of adult renal epithelial neoplasms.^21^ For the first time, we characterized the transcriptome of MA at the single-cell level. As shown in **Figure 6C** and D, hardly any CNVs were detected in the entire genome, and the CNV scores of MA are nearly equal to those of normal cells (NK/T cells). The differential genes were enriched in RNA metabolic process, ribosome biogenesis, and cytoplasmic translation expensive growth of MA in a benign way (**Figure 6G**). Due to lack of specificity in clinical presentation and imaging features. MA are often misdiagnosed. By comparing with ccRCC and chRCC cells, we discovered 10 candidate specific markers (*GPC3*, *ITM2C*, *PTGDS*, *COPG2*, *ENC1*, *EMX2*, *LHX1DT*, *BP1*, *DHRS2*, *DACH1*) of MA using FindeMarkers in Seurat (**Figure S8**). Overall, our study applied advanced single-cell technology to provide detailed and novel high-resolution insight into the heterogeneity of renal tumors. Multiple renal tumors, including RCCs and a benign tumor were characterized to better understand their molecular features. Our results show that ccRCC is related to hypoxia caused by *VHL* mutation; chRCC has a high degree of chromosomal instability and may have a higher proliferative capacity compared to the other two tumor samples; and MA is a benign renal neoplasm with minimal CNVs and differential genes enriched in benign processes. This knowledge may contribute to the development of more effective therapies for RCCs and the discovery of novel potential specific markers for renal tumors.

### Intercellular crosstalk in clear cell renal cell carcinomas

To comprehensively understand the complex microenvironment of malignant diseases, which contains a sophisticated mix of multiple cell types, the analysis of intercellular crosstalk has become increasingly important. Given that ccRCC accounts for approximately 80% of RCC cases^32^, cell-cell interactions in the ccRCC microenvironment plays a vital role in both tumor progression and treatment. As shown in **Figure 7A and B**, a total of 4645 single-cell transcriptome information was acquired, and 11 cell types were identified. To explore the cell-cell interaction with the ccRCC microenvironment, we applied CellChat^33^, a robust algorithm and a repository of known ligand-receptor interactions, for data analysis. We found that endothelial-endothelial cells have the largest ligand-receptor interaction pairs, followed by endothelial-CAF and endothelial-ccRCC (**Figure 7C, Figure S9**). Totally, CellChat detected 310 significant ligand-receptor pairs among the 9 cell types (ccRCC, ENDO, M2, NK/T cell, CAF, DC, M1, Proliferative T cell, and B cell), which were further categorized into 85 signaling pathways.

**Figure 7.**
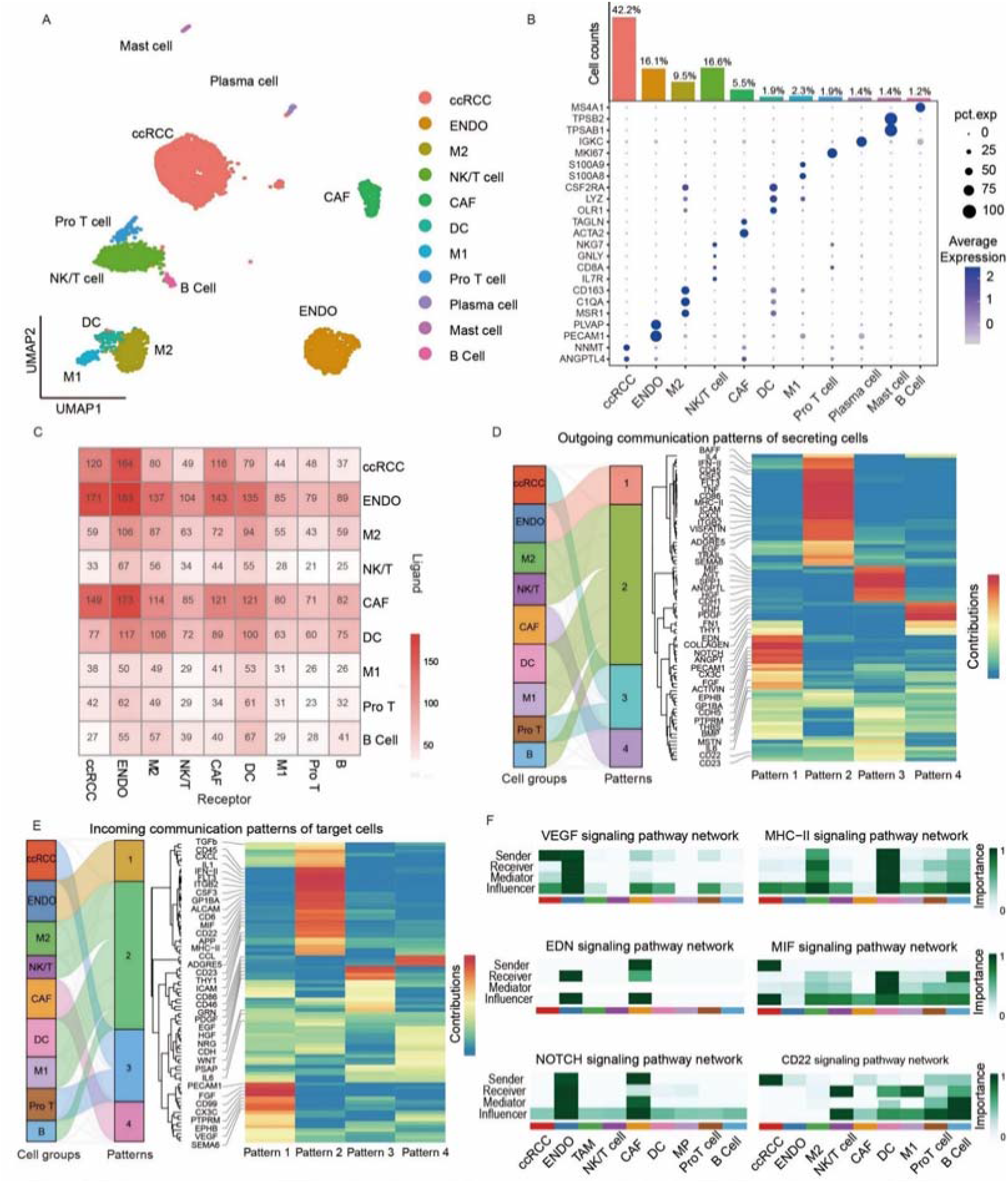
Cell-cell interaction in ccRCC single-cell transcription sample. A) Visualization by UMAP plot of clustering of 4645 single-cell expression profiles from the ccRCC sample: ccRCC, clear cell renal carcinoma cell; ENDO, endothelial cells; CAF, cancer-associated fibroblast; M2, macrophage cells 2; DC, dendritic cell; M1, macrophage cell1; Pro T cell, proliferative T cell. B) Histogram and dot plot showing the proportion of different cells in the ccRCC and gene expression patterns of cell-type-specific marker genes. C) CellChat showing significant ligand-receptor pairs among the 9 cell types. D, E) Outgoing and incoming communication patterns of target cells. F) Signaling pathway role heatmap showing indicated signaling pathway network in each cell type.

To characterize the detailed interaction for individual pathways, signaling patterns were displayed (**Figure 7D, E**). The pattern of outgoing and incoming signals revealed that ENDOs were highly enriched in pattern 1, representing multiple pathways, such as vascular endothelial growth factor (*VEGF*) and platelet endothelial cell adhesion molecule (*PECAM1*). Most immune cells, including M2, NK/T, DC, M1, and B cells, were mainly characterized by pattern 2, containing the pathways of major histocompatibility complex-II (*MHC-II*), epidermal growth factor (*EGF*), etc. ccRCC and proliferative T cells were significantly enriched in pattern 3, comprising migration inhibitory factor (*MIF*), cluster of differentiation-22 (*CD22*), cluster of differentiation-23 (*CD23*), etc. The last cell type CAF was prominently characterized by pattern 4, including endothelin (*EDN*), *NOTCH*, etc. It was the activation of the above adverse signaling pathways that prompted the progression of tumorigenesis, angiogenesis, and regulatory immune cell recruitment in the tumor microenvironment.

Therefore, we further explored the role of some signaling pathways in each cell type based on the pattern of outgoing and incoming signals. Consistent with the GO analysis (**Figure 6E**), the high enrichment in response to hypoxia and vascular process in the circulatory system, the *VEGF* signaling pathway role heatmap identified that ccRCC were the most prominent sources for *VEGF* ligands acting onto endothelial cells (**Figure 7F**), indicating that endothelial cells are stimulated to initiate angiogenesis by the larger oxygen demand of ccRCC tumor cells. In addition to the interaction between tumor cells and ENDO, it is worth noting that CAF, as the major components of the tumor stroma, also have a complex network of ligand-receptor interactions with the endothelium. Our analysis revealed that CAF are a significant source of *EDN* and *NOTCH* ligands (**Figure 7F**), which are crucial for the regulation of angiogenesis and promotion of tumor metastasis and progression.^34^

In addition, the highly vascularized ccRCC display high levels of immune cell infiltration (**Figure 7B**). There are numerous immune-associated processes enriched by GO analysis (**Figure 6E**). Consistently, various immune-associated signaling pathways were detected in pattern analysis. For instance, we found numerous *MHC-II* that were highly enriched in DCs, suggesting strong antigen presentation ability (**Figure 7F**). Moreover, strong interactions between tumor cells and immune cells mediated by *MIF, CD22*, and *CD23* pathways were observed (**Figure 7F**). These have been shown to specifically promote tumor angiogenesis, invasion, and immune evasion in ccRCC. For example, MIF has been found to enhance *VEGF*-induced angiogenesis and promote tumor growth^35^, while *CD22* and *CD23* can facilitate the immune escape of tumor cells by inhibiting immune cell activation and inducing regulatory T cell responses.^36^ Therefore, these pathways represent potential targets for the development of immunotherapeutic strategies for ccRCC. In summary, the use of WPS2 enables the comprehensive profiling of multiple pathologic transcriptome maps of clinical samples, which can aid in the identification of novel biomarkers and therapeutic targets for ccRCC and other cancers.

## Discussion

scRNA-seq have attracted extensive attentions for reveal the heterogenous cellular state. However, the throughput and sensitivity in scRNA-seq limit the ability to characterize the low RNA contents cells/nuclei in multicellular systems. Many efforts have been devoted to increase efficiency in single-cell capture and gene detection. Using WPS, we have successfully realized ∼100,000 single-cell analyses per flow channel. However, WPS still suffered from low sensitivity of gene detection. Well-Paired-Seq2 demonstrated high sensitivity in gene detection, benefiting from high efficiency in reverse transcription by molecular crowing effect and tailing activity enhancement and uniform amplification by homogeneous enzymatic reaction. These strategies could be readily extended to other scRNA-seq platforms. WPS2 identified 3116 median genes and 8447 transcripts at an average of 20,000 reads per cell, which is 1420 more genes and 4864 more transcripts compared to WPS. By adopting the new workflow of WPS2, we successfully applied it to characterize low RNA-content units, such as immune cells and nuclei, with high sensitivity and throughput.

In addition, the high cost and complicate peripheral equipment of current scRNA-seq techniques has hindered their accessibility. Well-Paired-Seq2 is a user-friendly platform. Compared to the high-cost and complicated 10x Genomics platform, the whole workflow of Well-Paired-Seq can be accomplished by a chip and pipette and only costs <US$0.05 per cell for library preparation, which can be established quickly and inexpensively in a standard biology lab. Moreover, it is crucial to note that the presence of cell-free RNA are significant challenges in snRNA-seq. a large number of RNAs in cytoplasm are released into solution in the process of nuclei extraction. Based on the locally quasi-hydrodynamic principle, a high effectiveness of removing these undesirable interfering substances can be achieved without cell loss, resulting in reduced background noise and a lower risk of generating inaccurate biological findings.

Next, we applied WPS2 to characterize clinical samples. We dissected the heterogenous CNVs of three subtypes and found that chRCC has more copy number variations (CNVs) in its genome, indicating the instability of the genome. The ccRCC has complex microenvironment and the highly expressed genes of ccRCC are associated with immune response, hypoxia, angiogenesis, etc. Notably, we profiled metanephric adenoma for the first time and some potential specific markers were revealed to faciliate accurate diagnosis of MA. In summary, the use of WPS2 enables profiling of multiple pathologic transcriptome maps of clinical samples, which can contribute to discover novel biomarkers and therapeutic targets for other cancers. we believe our high-sensitivity, high-throughput, and high-fidelity WPS2 platform that is compatible with scRNA and scRNA-seq, has great potential in cell biology, precision medicine, and reproductive biology.

## Methods

### Cell preparation

Mouse NIH 3T3 cells used in optimization experiments were obtained from the cell bank of the Chinese Academy of Sciences and were cultured in Dulbecco’s Modified Eagle Medium (DMEM, Thermo Fisher) supplemented with 10% fetal bovine serum (FBS, ThermoFisher) and 1% penicillin-streptomycin (Thermo Fisher) at 37 °C with 5% CO_2_. Cells were harvested by trypsinization and resuspended in cold Dulbecco’s Phosphate-Buffered Saline (DPBS, Corning).

The spleen tissue was obtained from C57BL/6J mice (JiangSu GemPharmatech Co., Ltd). The tissue was minced into 2-mm pieces on ice and rinsed with 1x DPBS, then treated with tissue dissociation mix (containing 1 mg/mL collagenase II and 5 mM CaCl_2_) for 15 min at 37 °C under rotation. The cell suspension was filtered through a 40-µm cell strainer and then centrifuged at 1200 rpm for 3 min at 4 °C. After the supernatant was removed, the cells were resuspended with red blood cell lysis buffer and incubated on ice for 3 min. The cell suspension was centrifuged, rinsed, and resuspended in Hanks. Cell count and viability were measured by the Automated cell counter (ThermoFisher).

The collection of kidney tumor tissues was approved by the Research Ethics Committee of Renji Hospital (KY2021-215-B), and informed consent was obtained from all patients. The single-cell suspensions of the clinic tumor tissues mentioned above were prepared by Tumor Dissociation Kit (Miltenyi Biotec).

### Nuclei preparation

NIH 3T3 cells were lysed with cold ATAC-Resuspension Buffer (RSB; 10 mM Tris-HCl, 10 mM NaCl and 3 mM MgCl_2_) containing 0.01% Digitonin, 0.1% NP40, and 0.1% Tween-20 (RSB-DNT) described for Omni-ATAC^37^. Tissue samples dissected from 6-8 weeks C57BL/6J mice or clinical samples were minced into 2-mm pieces, homogenized with a pre-chilled Dounce tissue grinder (Sigma, Cat #D8938) in 2 mL pre-cooled RSB-DNT (15 times with pastel A and 10 times with pastel B), and incubated on ice for 5 min with another 2 mL RSB-DNT. Nuclei were centrifuged at 500 g for 3 min at 4 °C. After centrifugation, the nuclei were washed twice and resuspended with RSB, then filtered through a 40-μm cell strainer, and diluted to a final concentration of 500 nuclei per mL for subsequent experiments.

### Barcoded bead preparation

Commercial barcoded beads were obtained from ChemGenes Company (Wilmington, Massachusetts, USA; cat. Macosko-2011-10 (V+)), as described in Drop-seq. The barcoded beads were washed twice with 30 mL of 100% ethanol and 30 mL of TE/TW (10 mM Tris pH 8.0, 1 mM EDTA, 0.01% Tween). Then the barcode beads were resuspended in 10 mL of TE/TW and placed at 4 °C until use. Before experiments, barcoded beads were resuspended in 20× TE (pH 8.0) and 50 mM DTT solution for subsequent capture in the chip.

### Well-Paired-Seq2 operation

For optimization experiments, the operation on the WPS2 chip was similar to that described for WPS. Compared with the WPS, WPS2 was designed with smaller cell-capture-wells to match the size of the nucleus. First, the 70 µL prepared nuclear suspension was introduced into the chip, and the nuclei were captured in the cell-capture-well by gravitational sedimentation. The uncaptured nuclei were resuspended with a pipette and carried through multiple sedimentation captures. The remaining uncaptured nuclei in the chip were removed with 1xDPBS, followed by loading the barcode beads were. The surfactant aggregates (sodium lauroyl sarcosine, Solarbio) were ultasonically dispersed evenly in mineral oil 5% (w/v). Then, the mineral oil containing the surfactant aggregates was injected into the chip and the dual wells were sealed. The surfactant aggregates settled and dissolved in the solution of the wells to complete cell lysis. After incubation at room temperature for 10 min, the barcode beads were recovered and transferred to a 1.5 mL RNase-free tube, washed three times with 1 mL of 6× SSC and once with 1× RT buffer.

### Reverse transcription and exonuclease I treatment

The barcoded beads were resuspended in 20 µL reverse transcription reaction solution, containing 1× RT buffer, 1mM dNTP (TransGen Biotech, cat#AD101-12), 1 U/µL RNase Inhibitor (TransGen Biotech, cat#AI101-02), 1 mM GTP (Thermo Scientific, cat#R0461), 5% PEG8000 (PERFEMIKER, cat# PMA020380), 2.5 μM Template Switch Oligo (AAGCAGTGGTATCAACGCAGAGTGAATrGrGrG, Sangon), and 10 U/µL Maxima H-Reverse Transcriptase (Thermo Scientific, cat#EP0751), and incubated at 42 °C for 90 min, followed by washing the beads once with 200 µL of 1× TE-SDS, once with 200 µL of 1× TE-TW, and once with 200 µL of 10 mM Tris (pH 8.0). The beads were then resuspended in 20 µL exonuclease I mix, containing 1× Exonuclease I Buffer and 1 U/µL Exonuclease I (NEB, cat # B0293S), and incubated at 37 °C for 45 min. After Exo I treatment, the beads were washed once with 200 µL of 1× TE-SDS, once with 200 µL of 1× TE-TW, and once with 200 µL of RNase free water.

### Second-strand synthesis on barcode beads

Following Exonuclease I treatment, the beads were resuspended in 200 µL of 0.1 M NaOH and incubated for 5 min at room temperature with rotation to denature the mRNA-cDNA hybrid product. After that, 200 µL of 10 mM Tris-HCl (pH 7.5) was added to neutralize the solution, the beads were then washed twice with 200 µL of TE-TW buffer and once with 200 µL of 10 mM Tris-HCl (pH 8.0). Second-strand synthesis reaction was performed on the beads by incubating in 40 µL of the reaction mixture (1x RT buffer, 12% PEG-8000, 1 mM dNTPs, 1 μM second strand synthesis primer (AAGCAGTGGTATCAACGCAGAGTGAATG, Sangon) and 0.125 U/μL Klenow exo^-^ (BioLabs)) at 37 °C for 1 h with rotation. The reaction was stopped by washing the beads once with TE-SDS buffer, twice with TE-TW buffer, and once with RNase-free water.

### cDNA amplification

The beads were resuspended in 50 µL PCR mix including 1× HiFi HotStart Readymix (Kapa Biosystems, cat #KK2602) and 0.8 μM ISPCR oligo (AAGCAGTGGTATCAACGCAGAGT, Sangon)). The PCR program was as follows: 95 °C for 3 min; four cycles of 98 °C for 20 s, 65 °C for 45 s, and 72 °C for 3 min; ten cycles of 98 °C for 20 s, 67 °C for 20 s, and 72 °C for 3 min; 72 °C for 5 min. The PCR product was then purified with 0.6× VAHTS DNA Clean Beads (Vazyme, cat#N411-02) according to the manufacturer’s instructions and eluted in 10 µL of H_2_O. The concentration of the purified PCR product was measured with a Qubit fluorometer (ThermoFisher Scientific).

### Well-Paired-Seq library preparation

The library was constructed with TruePrepDNA Library Prep Kit V2 for Illumina (Vazyme Biotech) according to the manufacturer’s instructions and was amplified by Nextera_N70x primer (Sangon) and custom primer P5_TSO_Hybrid (AATGATACGGCGACCACCGAGATCTACACGCCTGTCCGCGGAAGCAGTGG TATCAACGCAGAGT*A*C, Sangon). Then 0.6× VAHTS DNA Clean Beads were used for purifying the library. After elution in 10 µL of H_2_O, the concentration of the purified PCR production was measured with a Qubit fluorometer. The average size of sequenced libraries was between 450 and 750 bp.

## Statistical analysis

### Reads alignment and data preprocessing

The paired-end reads were produced from WPS2 sequencing libraries: read 1 contained a cell barcode at 1-12 bases and a UMI at 13-20 bases; read 2 contained the sequence of the transcript. GRCh38 and mm10 were used to align the human and mouse data, respectively. Data preprocessing and read alignment were implemented by using the zUMIs^38^. In order to correct the differences in sequencing depth between conditions, seqtk v1.0 (https://github.com/lh3/seqtk) was used to downsample the sequencing data, so that each condition could be analyzed nearly at the same level of average number of reads per cell. For samples sequenced with sc/snRNA-seq, each gene’s transcripts were counted by including exon and intron reads together.

### Cell type analysis

Based on digital gene expression, Seurat version 4.0.1^39^ was run on R4.1.3 to perform cell clustering and marker gene calling. The spleen cells and kidney nuclei were preprocessed and filtered on the basis of a minimal expression threshold of 300 and 500 genes and genes being expressed in at least three cells or nuclei. As high proportion of transcripts mapping to MT-genes indicate low cell quality, we removed cells with more than 30% MT-transcripts. Data normalization and scaling followed the suggested default settings and 2000 highest variable genes were selected using the FindVariableFeatures function with the vst method. Next, we performed PCA based on the scaled data and reduced the dimensionality of data based on the ElbowPlot. To cluster the cells, the FindClusters function in Seurat was implemented and visualized by projecting cells in a two-dimensional space using Uniform Manifold Approximation and Projection (UMAP), and cell types were identified by the cell marker. Similarly, the cell type analysis for the clinical samples data was implemented using cells with at least 500 detected genes and genes expressed in at least three cells. PCA was performed based on scaled data after the top 2000 variable genes were selected. Using the identified 20 PCs as input, cells were clustered into eight groups with resolution = 1.2, and the cell types were identified by the cell marker.

### DEGs and GO term enrichment

Differential expression analysis based on the wilcoxon rank sum test was then performed to confirm that the three tumor types were distinct. Genes with a fold change of transcripts >2 and an adjusted P value < 0.05 were recognized as differentially expressed genes. Furthermore, the type-specific pathways were revealed by Gene Ontology (GO) enrichment analysis for biological processes completed by the clusterProfiler R package.^40^

### Copy number variation analysis

The InferCNV package (https://github.com/broadinstitute/inferCNV) was used to detect the CNVs in 10704 malignant cells. 1623 non-malignant cells were used as baselines to estimate the CNAs of malignant cells. Genes expressed in more than 20 cells were sorted based on their loci on each chromosome. The relative expression values were centered to 1, using 1.5 standard deviations from the residual-normalized expression values as the ceiling. A slide window size of 101 genes was used to smooth the relative expression on each chromosome to remove the effect of gene-specific expression. The CNV scores were calculated by normalizing the CNVs of each cell from -1 to 1 and then calculating the sum of squares of the normalized values

### Cell-cell interactions based on CellChat^33^

Cell-cell interaction based on cell-chat begins with processing scRNA-seq data using standard quality control, normalization, and dimensionality reduction techniques. Then we used curated databases of known cell-cell signaling interactions to identify potential ligand-receptor interactions between cell types, which were then combined with expression data to create a communication matrix. This matrix was then used to cluster cell types into functional groups based on their signaling interactions, and the resulting communication network was visualized with default parameters.

## Supporting information

Table S1, and will be used for the link to the files on the preprint site

## Acknowledgements

We thank the National Key R&D Program of China (2021YFA0909400, 2019YFA0905800), the National Natural Science Foundation of China (21974113, 21974112, 21904085, and 21927806), and the Fundamental Research Funds for the Central Universities (20720210001, 20720220005) for their financial support.

## Data Availability Statement

The data that support the findings of this study are available from the corresponding author upon reasonable request.

## References

(1) Grün, D.; van Oudenaarden, A. Design and analysis of single-cell sequencing experiments. Cell 2015, 163 (4), 799–810.

(2) Tang, F.; Barbacioru, C.; Wang, Y.; Nordman, E.; Lee, C.; Xu, N.; Wang, X.; Bodeau, J.; Tuch, B. B.; Siddiqui, A. mRNA-Seq whole-transcriptome analysis of a single cell. Nat. Methods 2009, 6 (5), 377–382.

(3) Ramsköld, D.; Luo, S.; Wang, Y.-C.; Li, R.; Deng, Q.; Faridani, O. R.; Daniels, G. A.; Khrebtukova, I.; Loring, J. F.; Laurent, L. C. Full-length mRNA-Seq from single-cell levels of RNA and individual circulating tumor cells. Nat. Biotechnol. 2012, 30 (8), 777–782.

(4) Picelli, S.; Björklund, Å. K.; Faridani, O. R.; Sagasser, S.; Winberg, G.; Sandberg, R. Smart-seq2 for sensitive full-length transcriptome profiling in single cells. Nat. Methods 2013, 10 (11), 1096–1098.

(5) Hagemann-Jensen, M.; Ziegenhain, C.; Chen, P.; Ramsköld, D.; Hendriks, G.-J.; Larsson, A. J.; Faridani, O. R.; Sandberg, R. Single-cell RNA counting at allele and isoform resolution using Smart-seq3. Nat. Biotechnol. 2020, 38 (6), 708–714.

(6) Hashimshony, T.; Wagner, F.; Sher, N.; Yanai, I. CEL-Seq: single-cell RNA-Seq by multiplexed linear amplification. Cell Rep. 2012, 2 (3), 666–673.

(7) Hashimshony, T.; Senderovich, N.; Avital, G.; Klochendler, A.; De Leeuw, Y.; Anavy, L.; Gennert, D.; Li, S.; Livak, K. J.; Rozenblatt-Rosen, O. CEL-Seq2: sensitive highly-multiplexed single-cell RNA-Seq. Genome biology 2016, 17, 1–7.

(8) Sasagawa, Y.; Nikaido, I.; Hayashi, T.; Danno, H.; Uno, K. D.; Imai, T.; Ueda, H. R. Quartz-Seq: a highly reproducible and sensitive single-cell RNA sequencing method, reveals non-genetic gene-expression heterogeneity. Genome biology 2013, 14 (4), 1–17.

(9) Sasagawa, Y.; Danno, H.; Takada, H.; Ebisawa, M.; Tanaka, K.; Hayashi, T.; Kurisaki, A.; Nikaido, I. Quartz-Seq2: a high-throughput single-cell RNA-sequencing method that effectively uses limited sequence reads. Genome biology 2018, 19, 1–24.

(10) Macosko, E. Z.; Basu, A.; Satija, R.; Nemesh, J.; Shekhar, K.; Goldman, M.; Tirosh, I.; Bialas, A. R.; Kamitaki, N.; Martersteck, E. M. Highly parallel genome-wide expression profiling of individual cells using nanoliter droplets. Cell 2015, 161 (5), 1202–1214.

(11) McGinnis, C. S.; Patterson, D. M.; Winkler, J.; Conrad, D. N.; Hein, M. Y.; Srivastava, V.; Hu, J. L.; Murrow, L. M.; Weissman, J. S.; Werb, Z. MULTI-seq: sample multiplexing for single-cell RNA sequencing using lipid-tagged indices. Nat. Methods 2019, 16 (7), 619–626.

(12) Gierahn, T. M.; Wadsworth, M. H.; Hughes, T. K.; Bryson, B. D.; Butler, A.; Satija, R.; Fortune, S.; Love, J. C.; Shalek, A. K. Seq-Well: portable, low-cost RNA sequencing of single cells at high throughput. Nat. Methods 2017, 14 (4), 395–398. Datlinger, P.; Rendeiro, A. F.; Boenke, T.; Senekowitsch, M.; Krausgruber, T.; Barreca, D.; Bock, C. Ultra-high-throughput single-cell RNA sequencing and perturbation screening with combinatorial fluidic indexing. *Nat. Methods* 2021, *18* (6), 635-642.

(13) Han, L.; Wei, X.; Liu, C.; Volpe, G.; Zhuang, Z.; Zou, X.; Wang, Z.; Pan, T.; Yuan, Y.; Zhang, X. Cell transcriptomic atlas of the non-human primate Macaca fascicularis. Nature 2022, 604 (7907), 723–731.

(14) Consortium*, T. S.; Jones, R. C.; Karkanias, J.; Krasnow, M. A.; Pisco, A. O.; Quake, S. R.; Salzman, J.; Yosef, N.; Bulthaup, B.; Brown, P. The Tabula Sapiens: A multiple-organ, single-cell transcriptomic atlas of humans. Science 2022, 376 (6594), eabl4896. Suo, C.; Dann, E.; Goh, I.; Jardine, L.; Kleshchevnikov, V.; Park, J.-E.; Botting, R. A.; Stephenson, E.; Engelbert, J.; Tuong, Z. K. Mapping the developing human immune system across organs. *Science* 2022, *376* (6597), eabo0510.

(15) Yin, K.; Zhao, M.; Lin, L.; Chen, Y.; Huang, S.; Zhu, C.; Liang, X.; Lin, F.; Wei, H.; Zeng, H. WellLJPairedLJSeq: A SizeLJExclusion and Locally QuasiLJStatic Hydrodynamic Microwell Chip for SingleLJCell RNALJSeq. Small Methods 2022, 6 (7), 2200341.

(16) Bai, Z.; Deng, Y.; Kim, D.; Chen, Z.; Xiao, Y.; Fan, R. An integrated dielectrophoresis-trapping and nanowell transfer approach to enable double-sub-poisson single-cell RNA sequencing. ACS nano 2020, 14 (6), 7412–7424.

(17) Young, M. D.; Behjati, S. SoupX removes ambient RNA contamination from droplet-based single-cell RNA sequencing data. Gigascience 2020, 9 (12), giaa151.

(18) Habib, N.; Avraham-Davidi, I.; Basu, A.; Burks, T.; Shekhar, K.; Hofree, M.; Choudhury, S. R.; Aguet, F.; Gelfand, E.; Ardlie, K. Massively parallel single-nucleus RNA-seq with DroNc-seq. Nat. Methods 2017, 14 (10), 955–958.

(19) Slyper, M.; Porter, C. B.; Ashenberg, O.; Waldman, J.; Drokhlyansky, E.; Wakiro, I.; Smillie, C.; Smith-Rosario, G.; Wu, J.; Dionne, D. A single-cell and single-nucleus RNA-Seq toolbox for fresh and frozen human tumors. Nat. Med. 2020, 26 (5), 792–802.

(20) Ding, J.; Adiconis, X.; Simmons, S. K.; Kowalczyk, M. S.; Hession, C. C.; Marjanovic, N. D.; Hughes, T. K.; Wadsworth, M. H.; Burks, T.; Nguyen, L. T. Systematic comparison of single-cell and single-nucleus RNA-sequencing methods. Nat. Biotechnol. 2020, 38 (6), 737–746.

(21) Davis Jr, C. J.; Barton, J. H.; Sesterhenn, I. A.; Mostofi, F. Metanephric adenoma. Clinicopathological study of fifty patients. The American journal of surgical pathology 1995, 19 (10), 1101–1114. Spaner, S. J.; Yu, Y.; Cook, A. J.; Boag, G. Pediatric metanephric adenoma: case report and review of the literature. *International urology and nephrology* 2014, *46*, 677-680.

(22) Ohtsubo, Y.; Nagata, Y.; Tsuda, M. Compounds that enhance the tailing activity of Moloney murine leukemia virus reverse transcriptase. Scientific reports 2017, 7 (1), 6520.

(23) Bagnoli, J. W.; Ziegenhain, C.; Janjic, A.; Wange, L. E.; Vieth, B.; Parekh, S.; Geuder, J.; Hellmann, I.; Enard, W. Sensitive and powerful single-cell RNA sequencing using mcSCRB-seq. Nature communications 2018, 9 (1), 2937.

(24) Hrdlickova, R.; Toloue, M.; Tian, B. RNALJSeq methods for transcriptome analysis. Wiley Interdisciplinary Reviews: RNA 2017, 8 (1), e1364.

(25) Griffiths, J. A.; Scialdone, A.; Marioni, J. C. Using singleLJcell genomics to understand developmental processes and cell fate decisions. Mol. Syst. Biol. 2018, 14 (4), e8046.

(26) Mazutis, L.; Gilbert, J.; Ung, W. L.; Weitz, D. A.; Griffiths, A. D.; Heyman, J. A. Single-cell analysis and sorting using droplet-based microfluidics. Nat. Protoc. 2013, 8 (5), 870–891.

(27) Su, C.; Lv, Y.; Lu, W.; Yu, Z.; Ye, Y.; Guo, B.; Liu, D.; Yan, H.; Li, T.; Zhang, Q. Single-cell RNA sequencing in multiple pathologic types of renal cell carcinoma revealed novel potential tumor-specific markers. Frontiers in Oncology 2021, 11, 719564.

(28) Schreibing, F.; Kramann, R. Mapping the human kidney using single-cell genomics. Nature Reviews Nephrology 2022, 18 (6), 347–360.

(29) Clark, D. J.; Dhanasekaran, S. M.; Petralia, F.; Pan, J.; Song, X.; Hu, Y.; da Veiga Leprevost, F.; Reva, B.; Lih, T.-S. M.; Chang, H.-Y. Integrated proteogenomic characterization of clear cell renal cell carcinoma. Cell 2019, 179 (4), 964–983. e931.

(30) Ohe, C.; Kuroda, N.; Takasu, K.; Senzaki, H.; Shikata, N.; Yamaguchi, T.; Miyasaka, C.; Nakano, Y.; Sakaida, N.; Uemura, Y. Utility of immunohistochemical analysis of KAI1, epithelial-specific antigen, and epithelial-related antigen for distinction of chromophobe renal cell carcinoma, an eosinophilic variant from renal oncocytoma. Medical molecular morphology 2012, 45, 98–104.

(31) Chen, C. V.; Croom, N. A.; Simko, J. P.; Stohr, B. A.; Chan, E. Differential immunohistochemical and molecular profiling of conventional and aggressive components of chromophobe renal cell carcinoma: pitfalls for diagnosis. Human Pathology 2022, 119, 85–93.

(32) Shuch, B.; Amin, A.; Armstrong, A. J.; Eble, J. N.; Ficarra, V.; Lopez-Beltran, A.; Martignoni, G.; Rini, B. I.; Kutikov, A. Understanding pathologic variants of renal cell carcinoma: distilling therapeutic opportunities from biologic complexity. Eur. Urol. 2015, 67 (1), 85–97.

(33) Jin, S.; Guerrero-Juarez, C. F.; Zhang, L.; Chang, I.; Ramos, R.; Kuan, C.-H.; Myung, P.; Plikus, M. V.; Nie, Q. Inference and analysis of cell-cell communication using CellChat. Nature communications 2021, 12 (1), 1088.

(34) Rio, D. D.; Caprara, V.; Masi, I.; Spadaro, F.; Giannitelli, S.; Rainer, A.; Bagnato, A.; Rosanò, L. Tumor-derived endothelin-1 recruits and activates fibroblasts to support tumor aggressiveness. Cancer Res. 2022, 82 (12_Supplement), 6137–6137. Katarkar, A.; Bottoni, G.; Clocchiatti, A.; Goruppi, S.; Bordignon, P.; Lazzaroni, F.; Gregnanin, I.; Ostano, P.; Neel, V.; Dotto, G. P. NOTCH1 gene amplification promotes expansion of Cancer Associated Fibroblast populations in human skin. *Nature communications* 2020, *11* (1), 5126.

(35) Noe, J. T.; Mitchell, R. A. MIF-dependent control of tumor immunity. Frontiers in immunology 2020, 11, 609948. Hu, H.; Ma, T.; Liu, N.; Hong, H.; Yu, L.; Lyu, D.; Meng, X.; Wang, B.; Jiang, X. Immunotherapy checkpoints in ovarian cancer vasculogenic mimicry: Tumor immune microenvironments, and drugs. *Int. Immunopharmacol.* 2022, *111*, 109116.

(36) Clark, E. A.; Giltiay, N. V. CD22: a regulator of innate and adaptive B cell responses and autoimmunity. Frontiers in immunology 2018, 9, 2235. Engeroff, P.; Vogel, M. The role of CD23 in the regulation of allergic responses. *Allergy* 2021, *76* (7), 1981-1989.

(37) Corces, M. R.; Trevino, A. E.; Hamilton, E. G.; Greenside, P. G.; Sinnott-Armstrong, N. A.; Vesuna, S.; Satpathy, A. T.; Rubin, A. J.; Montine, K. S.; Wu, B. An improved ATAC-seq protocol reduces background and enables interrogation of frozen tissues. Nat. Methods 2017, 14 (10), 959–962.

(38) Parekh, S.; Ziegenhain, C.; Vieth, B.; Enard, W.; Hellmann, I. zUMIs-A fast and flexible pipeline to process RNA sequencing data with UMIs. Gigascience 2018.

(39) Stuart, T.; Butler, A.; Hoffman, P.; Hafemeister, C.; Papalexi, E.; Mauck, W. M.; Hao, Y.; Stoeckius, M.; Smibert, P.; Satija, R. Comprehensive integration of single-cell data. Cell 2019, 177 (7), 1888–1902. e1821.

(40) Yu, G.; Wang, L.-G.; Han, Y.; He, Q.-Y. clusterProfiler: an R package for comparing biological themes among gene clusters. OMICS: J. Integrative Biol. 2012, 16 (5), 284–287.

