## Supplementary material for "Well-Paired-Seq2: High-Throughput and High-Sensitivity Strategy for Characterizing Low RNA-Content Cell/Nucleus Transcriptomes": Table S1, and will be used for the link to the files on the preprint site

Well-Paired-Seq2 Overcomes the Poisson Limit and Characterizes Transcriptomes of Low RNA-content Cells/Nuclei with High Sensitivity

Kun Yin, Meijuan Zhao, Yiling Xu, Zhong Zheng, Shanqing Huang, Dianyi Liang, He Dong, Ye Guo, Li Lin, Jia Song, Huiming Zhang, Junhua Zheng, Zhi Zhu, Chaoyong Yang

These authors contributed equally: Kun Yin, Meijuan Zhao, Yiling Xu, Zhong Zheng.

Kun Yin, Meijuan Zhao, Yiling Xu, Shanqing Huang, Dianyi Liang, He Dong, Ye Guo, Li Lin, Zhi Zhu, Chaoyong Yang

State Key Laboratory of Physical Chemistry of Solid Surfaces,

The MOE Key Laboratory of Spectrochemical Analysis & Instrumentation

Key Laboratory for Chemical Biology of Fujian Province

Collaborative Innovation Center of Chemistry for Energy Materials, Department of Chemical Biology, College of Chemistry and Chemical Engineering

Xiamen University

Xiamen 361005, P. R. China

Zhong Zheng, Jia Song, Junhua Zheng, Chaoyong Yang

Institute of Molecular Medicine

State Key Laboratory of Oncogenes and Related Genes

Renji Hospital, School of Medicine

Shanghai Jiao Tong University

Shanghai, 200120,China

Huiming Zhang, Chaoyong Yang

Innovation Laboratory for Sciences and Technologies of Energy Materials of Fujian Province (IKKEM)

Xiamen 361005, P. R. China

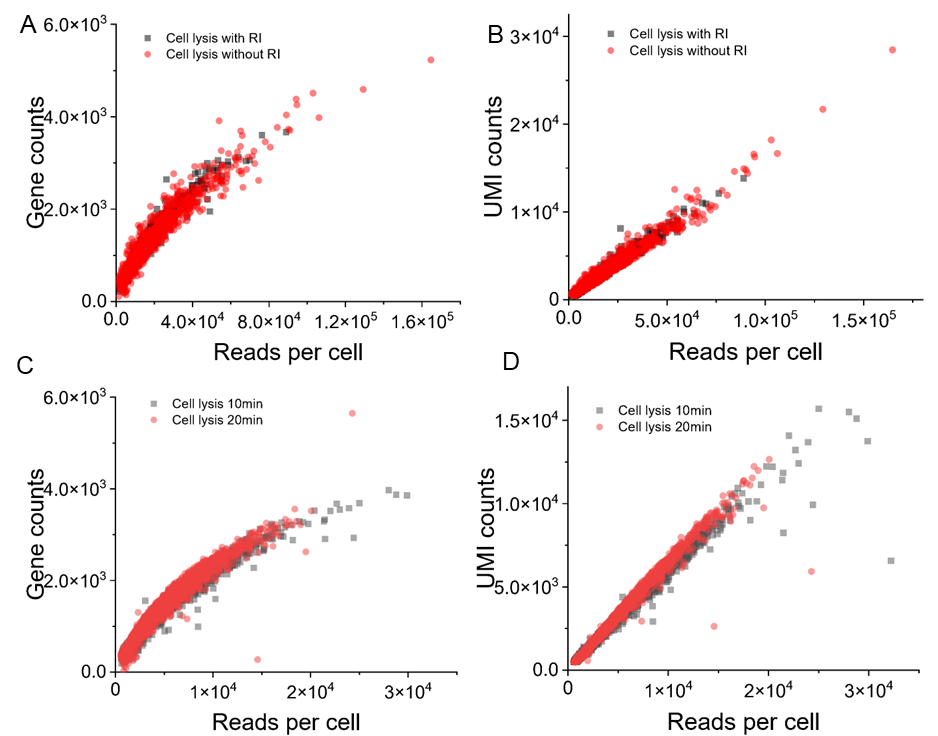
 Figure S1. Optimization of cell lysis conditions based on WPS. A, B Scatter plot showing the gene count and transcript number versus the read number of each individual cell for different cell lysis times (10 min and 20 min). C, D Different cell lysis buffers (with or without RNase inhibitor).

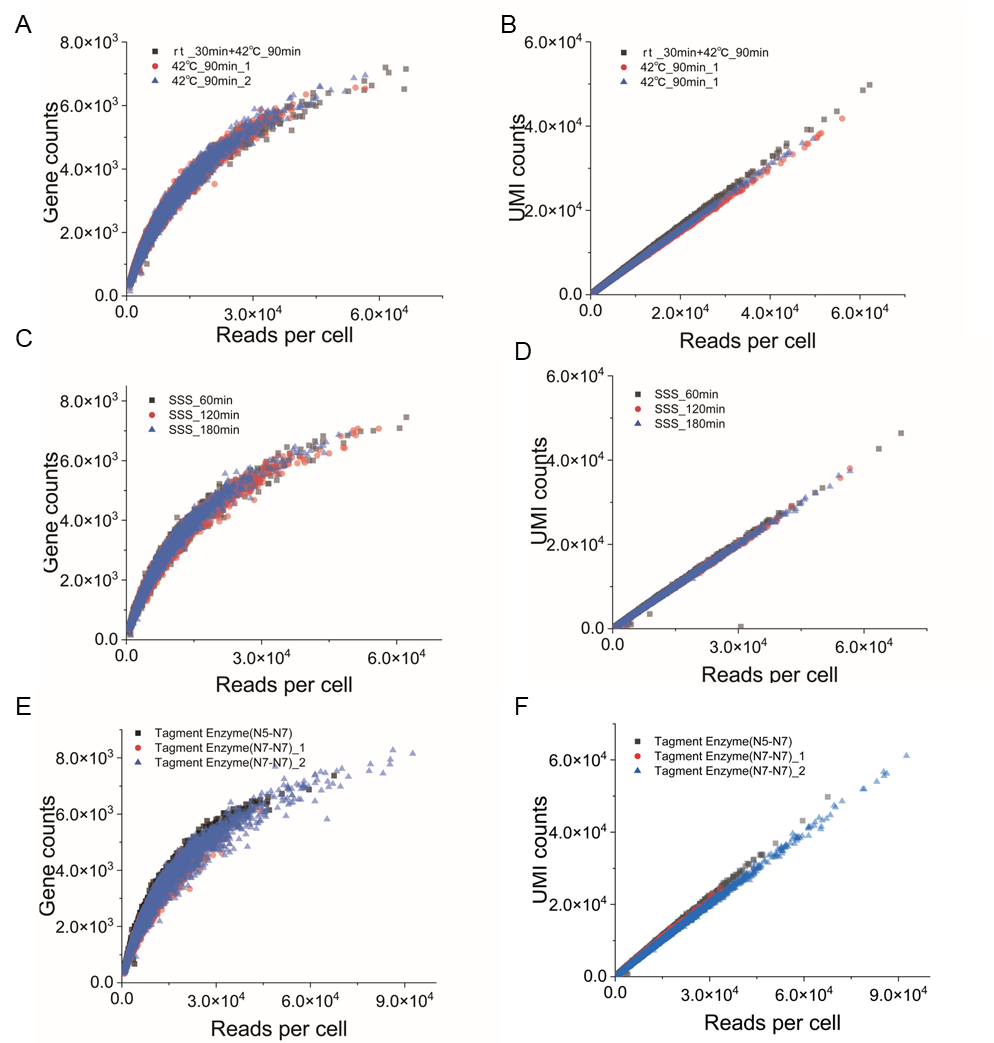
 Figure S2. Optimization of the processes after capturing mRNA based on WPS2. A, B. Detection of gene and UMIs in different reverse transcription conditions (with or without incubation at room temperature 30min before 42℃ 90 min). C, D. Second-strand synthesis time (60min, 120min, 180min). E, F. Tagment enzyme used in tagmentation (N5-N7adapter transposome complex and N7 single-adapter transposome complex).

A
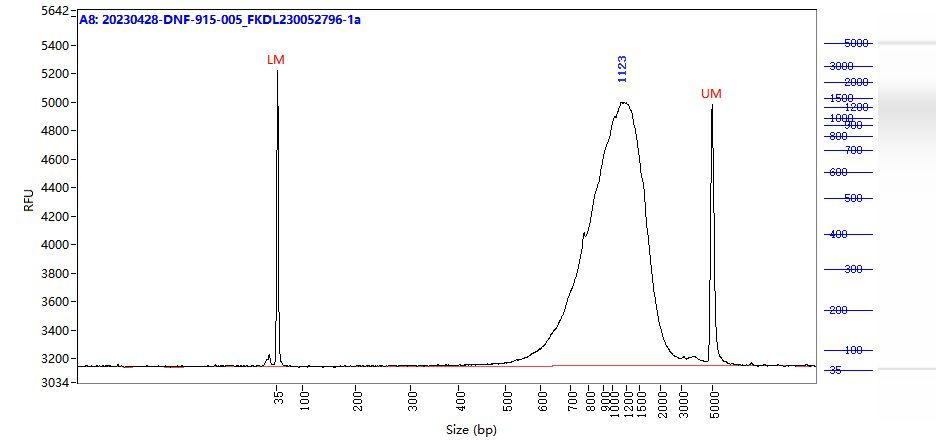

B
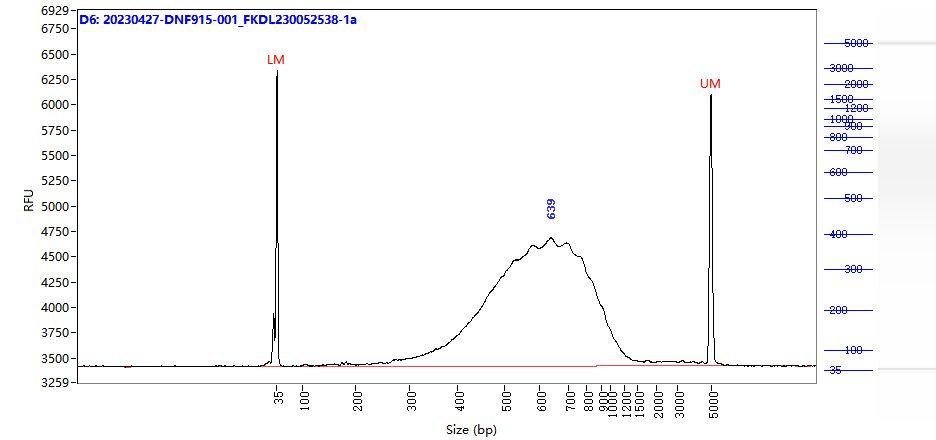

Figure S3. Length distribution of cDNA with RGP buffer（A）and RGPMB buffer (B). Length distribution of cDNA with RGP buffer is more expected than with RGPMB buffer.

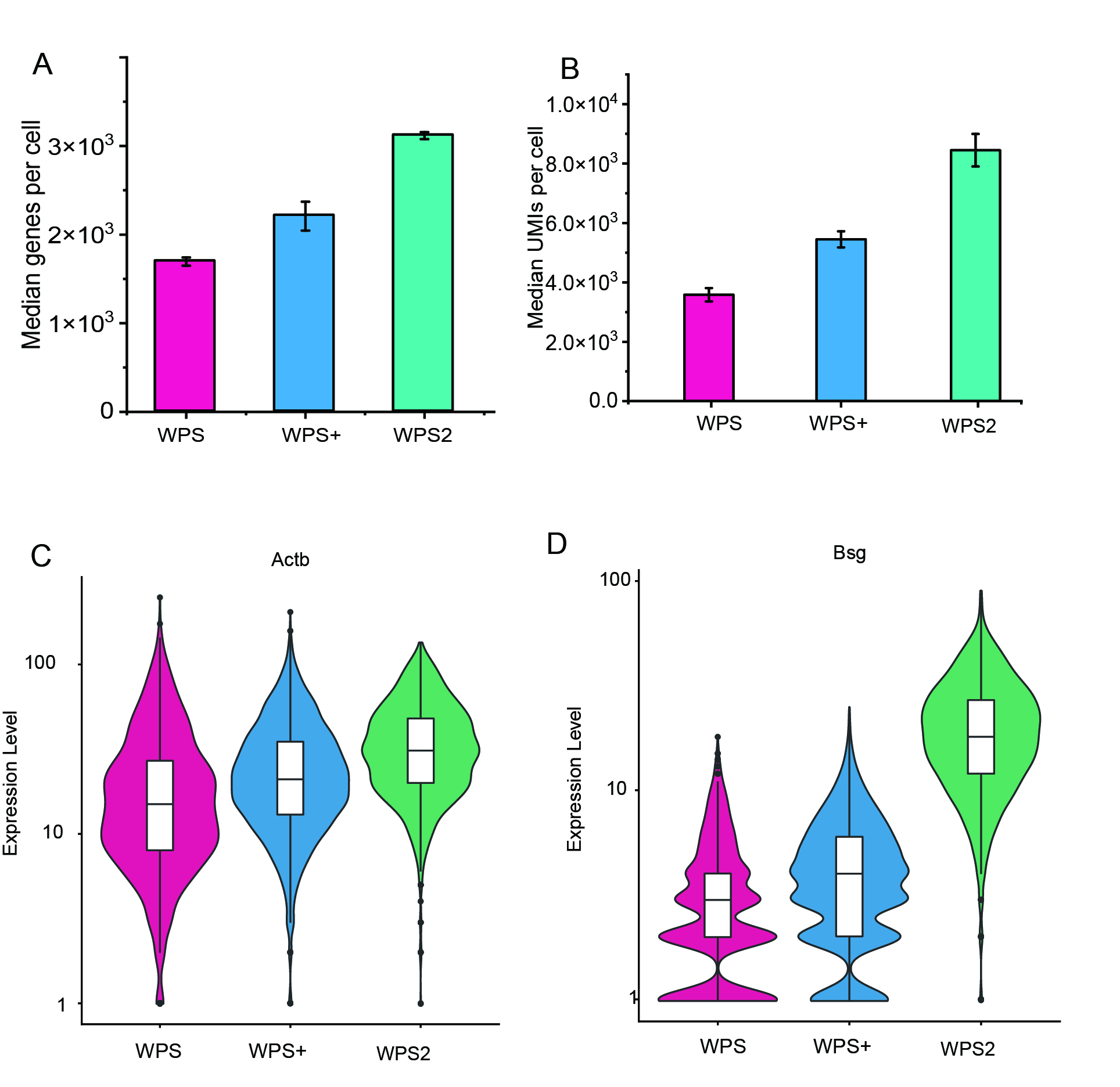

Figure S4. Performance validation of WPS 2. A, B. Median gene and UMI detection at an average of 20,000 reads per cell. C, D. Violin plots showing the expression level of given genes (housekeeping genes: Actb; variation genes: Bsg) detected by WPS, WPS+, and WPS2.

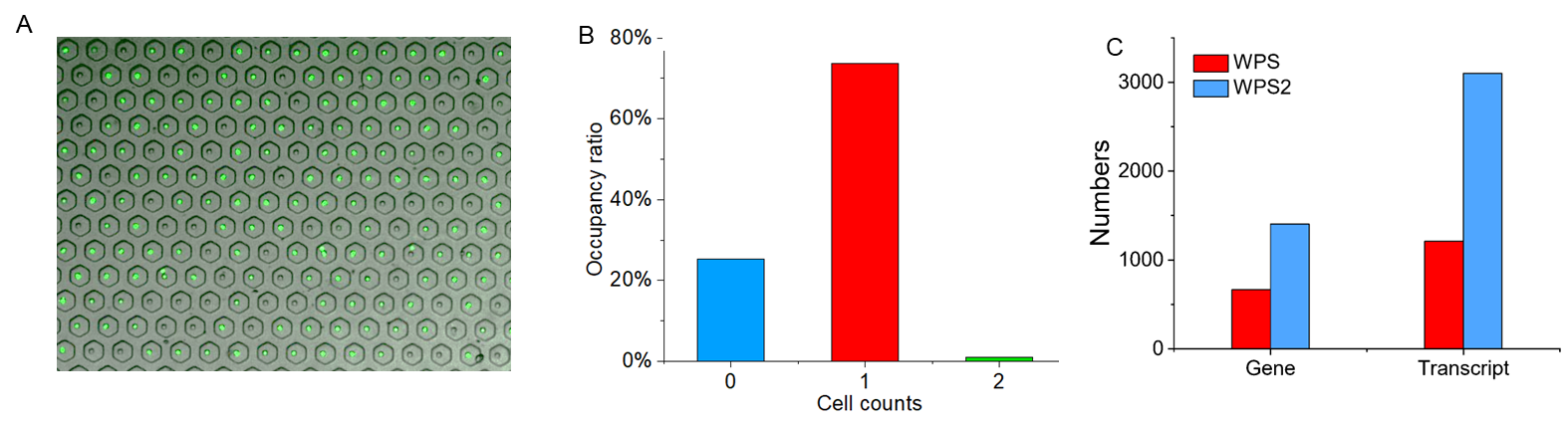
 Figure S5. Performance of the WPS2. A. Image of the WPS2 chip after cell capture. B. Statistical chart of cell occupancy ratio before recovery. C. Comparison of the average number of genes and transcripts between methods.

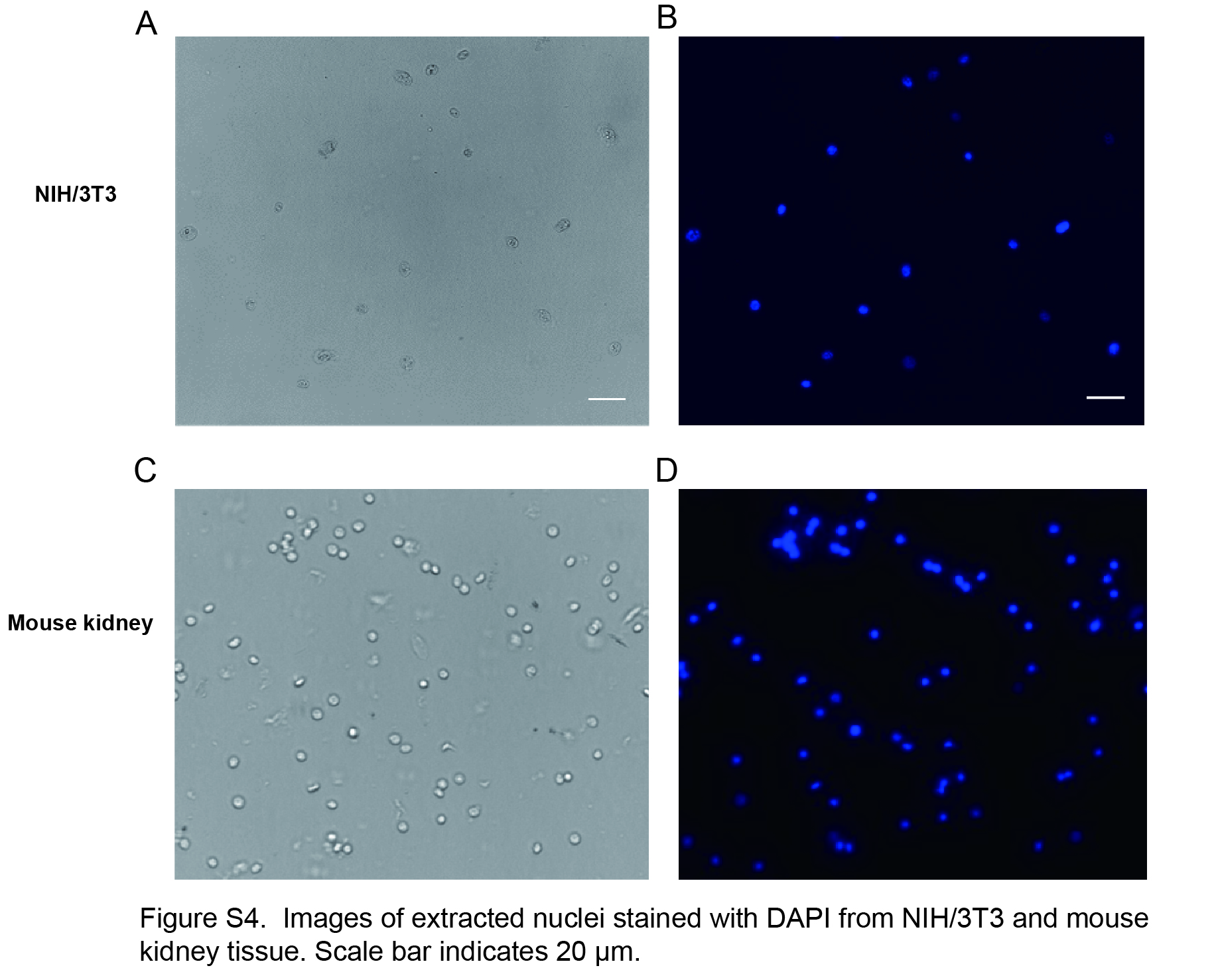
 Figure S6. Images of extracted nuclei stained with DAPI from NIH/3T3 and mouse kidney tissue. Scale bar indicates 20 µm.

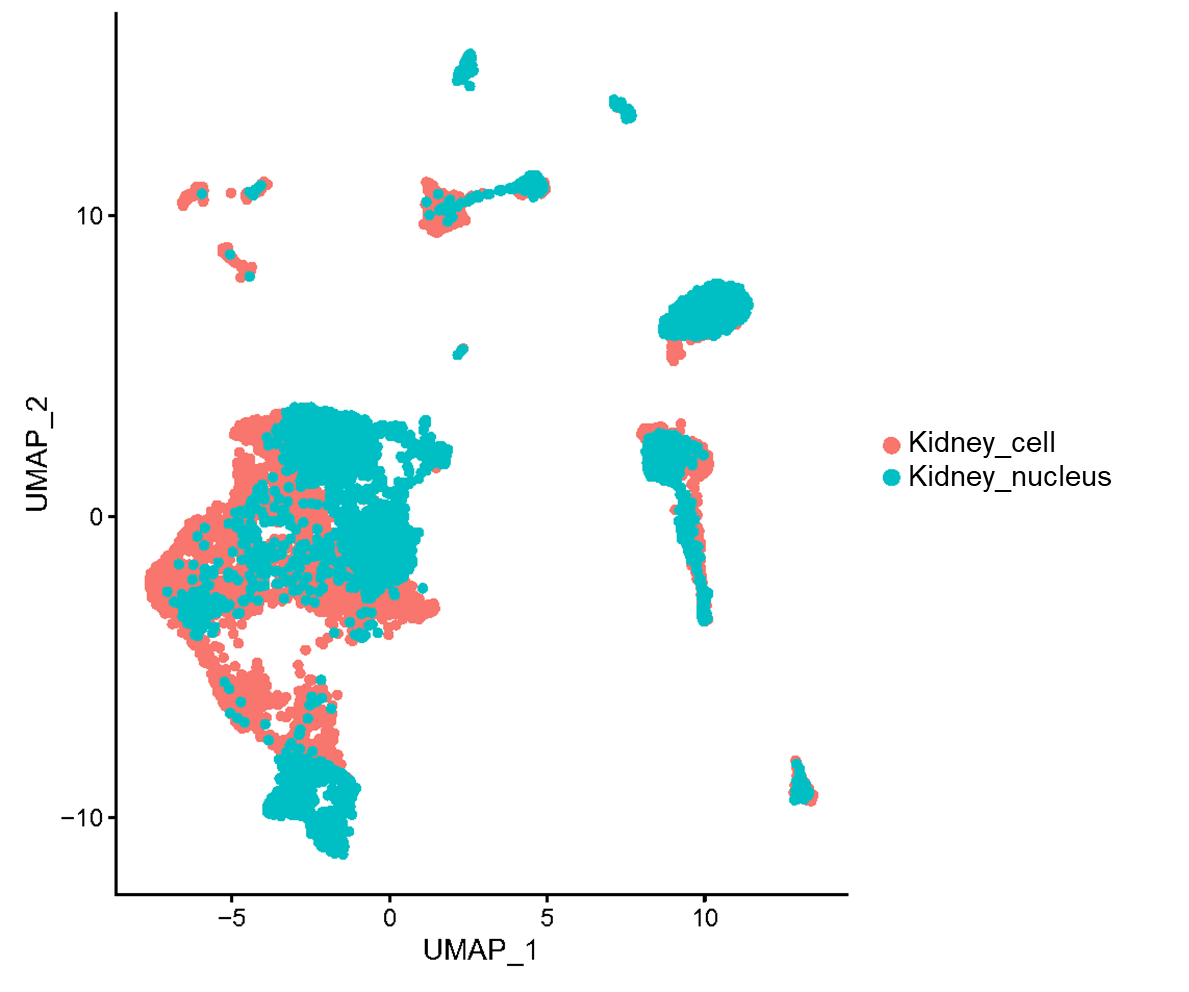
Figure S7. UMAP plot showing the correlation between scRNA-seq and snRNA-seq data from mouse kisney tissue.

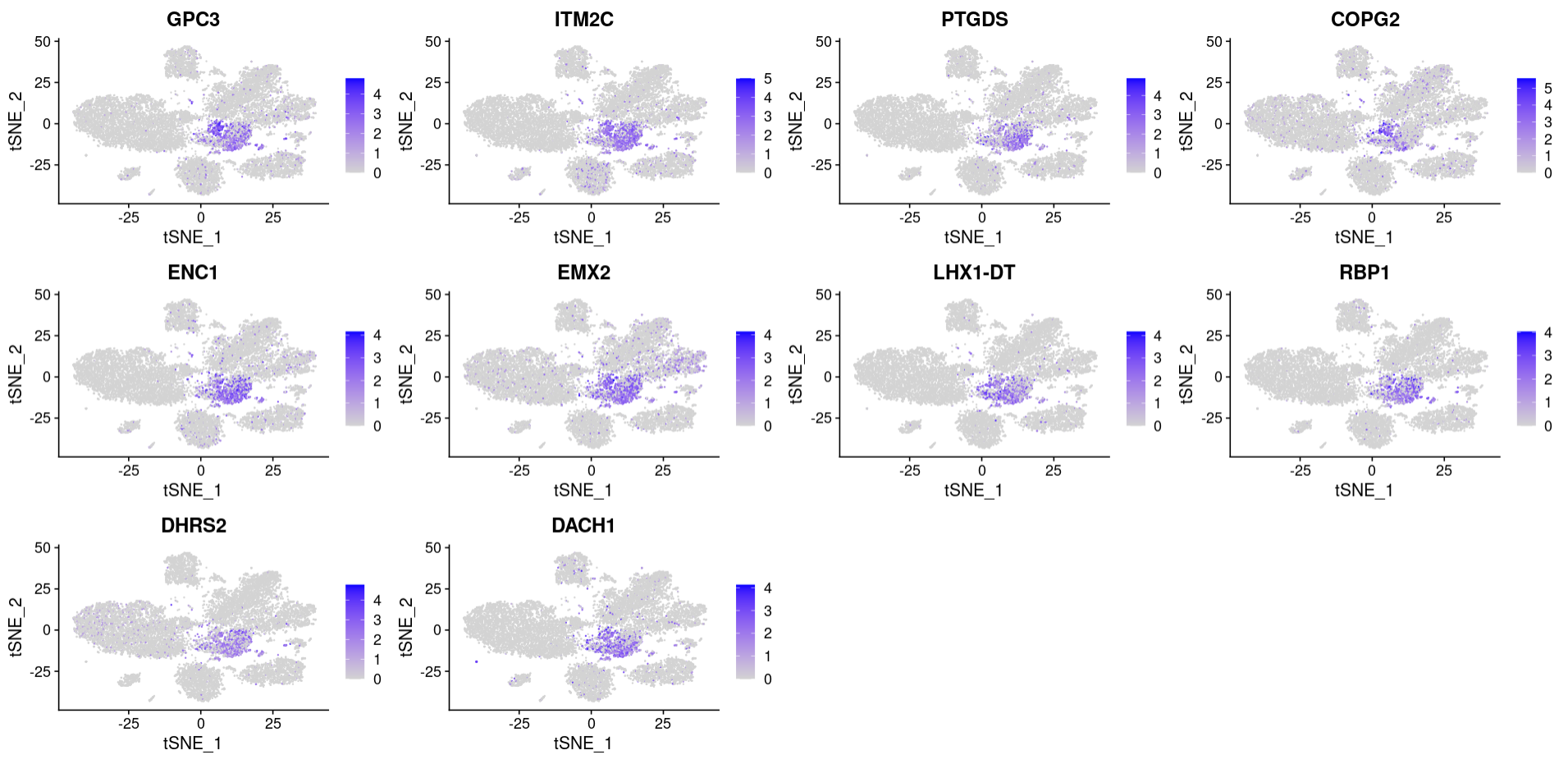
Figure S8. Candidate-specific markers of metanephric adenoma (MA).

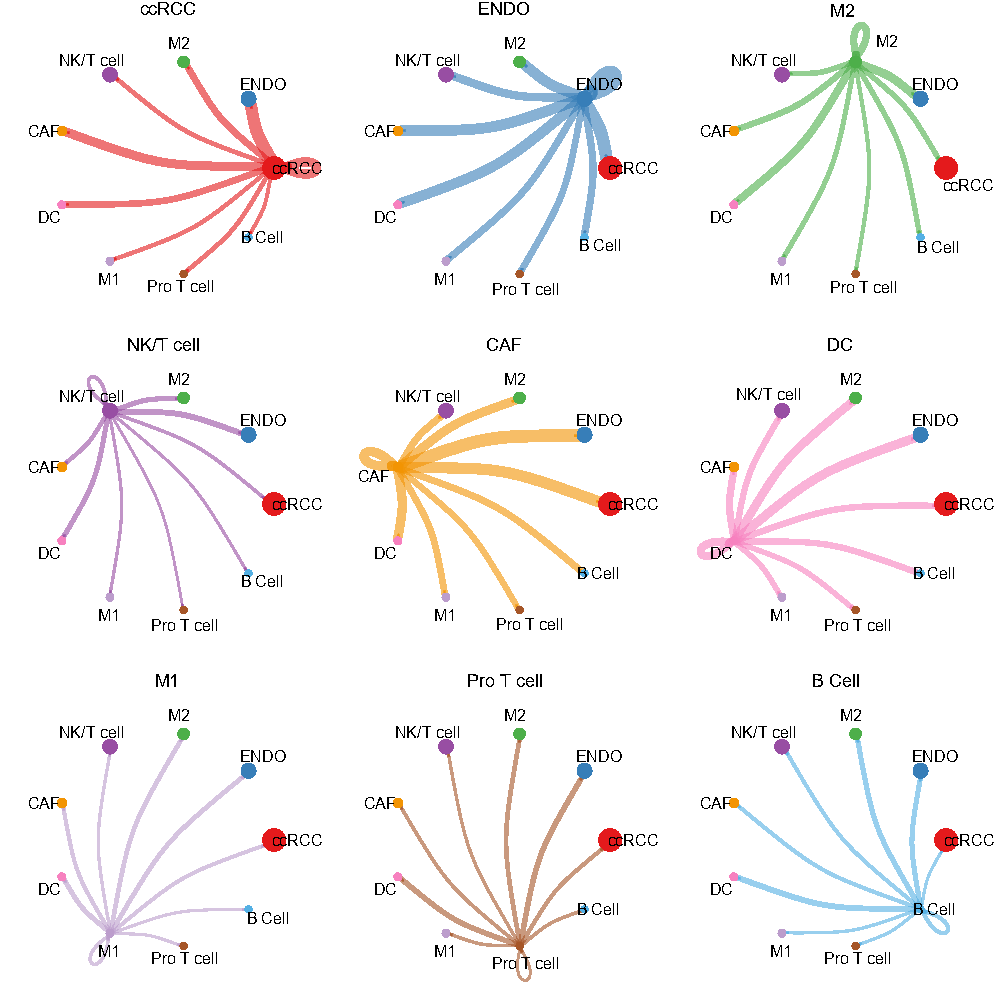
Figure S9. Cell–cell interaction network in the ccRCC microenvironment.

| Cell types | Genes | Transcripts |
| --- | --- | --- |
| TAL (thick ascending limb of Henle's loop cell) | 1171 | 2177 |
| PT_S1 (proximal tubule segment 1) | 1310 | 2457 |
| PT_S2 (proximal tubule segment 2) | 1258 | 2427 |
| PT_S3 (proximal tubule segment 3) | 1675 | 3398 |
| DCT (distal convoluted tubule cell) | 1022 | 1842 |
| CNT (connecting tubule cell) | 1210 | 2265 |
| ENDO (endothelial cell) | 884 | 1409 |
| MC (mesangial cell) | 1036 | 1763 |
| IC (intercalated cell) | 1100 | 1947 |
| DTL (descending thin limb of Henle's loop cell) | 1091 | 1955 |
| PC (principal cell) | 969 | 1617 |
| PODO (podocyte) | 1147 | 2227 |
| MACRO (macrophage cell) | 1036 | 1707 |
| Average | 1309 | 2329 |

Table S1. The number of genes and transcripts detected in each cell type

| Abbreviations | Full names |
| --- | --- |
| RGP buffer | Maxima H Minus RT buffer, GTP, PEG |
| RGPMB buffer | Maxima H Minus RT buffer, GTP, PEG, Mg^2+^, betaine |
| WPS2 | Well-Paired-Seq2 |
| WPS | Well-Paired-Seq |
| WPS+ | WPS with RGP buffer |
| RCCs | renal cell carcinomas |
| ccRCC | clear cell renal cell carcinoma |
| chRCC | chromophobe renal cell carcinoma |
| MA | metanephric adenoma |
| Mono progenitor | monocyte progenitor |
| DC | Dendritic cell |
| TAL | thick ascending limb of Henle's loop cell |
| PT_S1 | proximal tubule segment 1 |
| PT_S2 | proximal tubule segment 2 |
| PT_S3 | proximal tubule segment 3 |
| DCT | distal convoluted tubule cell |
| CNT | connecting tubule cell |
| ENDO | endothelial cell |
| MC | mesangial cell |
| IC | intercalated cell |
| DTL | descending thin limb of Henle's loop cell |
| PC | principal cell |
| PODO | podocyte |
| MACRO | macrophage cell |
| M1 | macrophage cell 1 |
| M2 | macrophage cell 2 |
| Pro T cell | proliferative T Cell |
| CAF | cancer-associated fibroblast |
| VEGF | vascular endothelial growth factor |
| PECAM1 | platelet endothelial cell adhesion molecule |
| MHC-II | major histocompatibility complex-II |
| EGF | epidermal growth factor |
| MIF | migration inhibitory factor |
| CD22 | cluster of differentiation-22 |
| CD23 | cluster of differentiation-23 |
| EDN | endothelin |
| VHL | Von Hippel-Lindau |

Table S2. The list of Abbreviations and its corresponding full names
